# The Calpain-7 protease functions together with the ESCRT-III protein IST1 within the midbody to regulate the timing and completion of abscission

**DOI:** 10.1101/2022.10.18.512775

**Authors:** Elliott L. Paine, Jack J. Skalicky, Frank G. Whitby, Douglas R. Mackay, Katharine S. Ullman, Christopher P. Hill, Wesley I. Sundquist

## Abstract

The Endosomal Sorting Complexes Required for Transport (ESCRT) machinery mediates the membrane fission step that completes cytokinetic abscission and separates dividing cells. Filaments composed of ESCRT-III subunits constrict membranes of the intercellular bridge midbody to the abscission point. These filaments also bind and recruit cofactors whose activities help execute abscission and/or delay abscission timing in response to mitotic errors via the NoCut/Abscission checkpoint. We previously showed that the ESCRT-III subunit IST1 binds the cysteine protease CAPN7 (Calpain-7) and that CAPN7 is required for both efficient abscission and NoCut checkpoint maintenance (Wenzel *et al*., 2022). Here, we report biochemical and crystallographic studies showing that the tandem MIT domains of CAPN7 bind simultaneously to two distinct IST1 MIT interaction motifs. Structure-guided point mutations in either CAPN7 MIT domain disrupted IST1 binding in vitro and in cells, and depletion/rescue experiments showed that the CAPN7-IST1 interaction is required for: 1) CAPN7 recruitment to midbodies, 2) efficient abscission, and 3) NoCut checkpoint arrest. CAPN7 proteolytic activity is also required for abscission and checkpoint maintenance. Hence, IST1 recruits CAPN7 to midbodies, where its proteolytic activity is required to regulate and complete abscission.

## Introduction

Midbody abscission separates two dividing cells at the end of cytokinesis. The ESCRT pathway is central to abscission and its regulation via the NoCut/Abscission checkpoint, whereby abscission delay allows mitotic errors to be resolved (Carlton and Martin-Serrano, 2007; Morita *et al*., 2007; Carlton *et al*., 2012; Capalbo *et al*., 2012; Capalbo *et al*., 2016; Scourfield and Martin-Serrano, 2017). Humans express 12 distinct ESCRT-III proteins that are recruited to membrane fission sites throughout the cell, including to midbodies that connect dividing cells during cytokinesis. Within the midbody, ESCRT-III proteins copolymerize into filaments that constrict the membrane to the fission point (Guizetti *et al*., 2011; Elia *et al*., 2011; Mierzwa *et al*., 2017; Nguyen *et al*., 2020; Pfitzner *et al*., 2021, Azad *et al*., 2022). ESCRT-III filaments also recruit a variety of cofactors, including the VPS4 AAA+ ATPases, which dynamically remodel the filaments to drive midbody constriction (Elia *et al*., 2012; Mierzwa *et al*., 2017; Pfitzner *et al*., 2020; Pfitzner *et al*., 2021). Many of these cofactors contain MIT domains, which bind differentially to MIT-Interacting Motifs (MIMs) located near the C-termini of the different human ESCRT-III proteins (Hurley and Yang, 2008; Wenzel *et al*., 2022).

Our recent quantitative survey of the human ESCRT-III-MIT interactome revealed a series of novel interactions between ESCRT-III subunits and MIT cofactors and implicated a subset of these cofactors in abscission and NoCut checkpoint maintenance (Wenzel *et al*., 2022). One such cofactor was CAPN7 (Calpain-7), a ubiquitously expressed but poorly understood cysteine protease that had not previously been linked to abscission nor to the NoCut checkpoint. CAPN7 contains tandem MIT domains that can bind specifically to the ESCRT-III subunit IST1 (Osako *et al*., 2010; Wenzel *et al*., 2022). This interaction can activate CAPN7 autolysis and proteolytic activity towards non-physiological substrates (Osako *et al*., 2010; Maemoto *et al*., 2013), although authentic CAPN7 substrates are not yet known. We undertook the current studies with the goals of defining precisely how CAPN7 and IST1 interact, how CAPN7 is recruited to midbodies, and whether CAPN7 must function as a midbody protease in order to support efficient abscission and maintain NoCut checkpoint signaling.

## Results and Discussion

### The CAPN7 MIT domains bind distinct IST1 MIM elements

The tandem CAPN7 MIT domains (CAPN7(MIT)2) bind a C-terminal region of IST1 that contains two distinct MIM elements (Figure 1A) (Agromayor *et al*., 2009; Bajorek *et al*., 2009; Osako *et al*., 2010; Wenzel *et al*., 2022). We quantified this interaction using fluorescence polarization anisotropy binding assays with purified, recombinant CAPN7(MIT)_2_ and fluorescently labeled IST constructs that spanned either one or both of the MIM elements. These experiments showed that both IST1 MIMs contribute to binding (Figure 1B) (Osako *et al*., 2010, Wenzel *et al*., 2022). The double MIM IST1_316-366_ construct bound CAPN7(MIT)_2_ tightly (K_D_ = 0.09 ± 0.01 μM), and the first IST1 MIM element (IST1344-366) contributed most of the binding energy (K_D_ = 1.8 ± 0.1 μM). The second IST1 MIM element (MIM_316-343_) bound very weakly in isolation (K_D_ > 100 μM), but CAPN7(MIT)_2_ binding affinity was reduced by more than 20-fold when this element was removed, indicating that both MIM elements contribute to binding, in good agreement with previous measurements (Wenzel *et al*., 2022).

**Figure 1.**
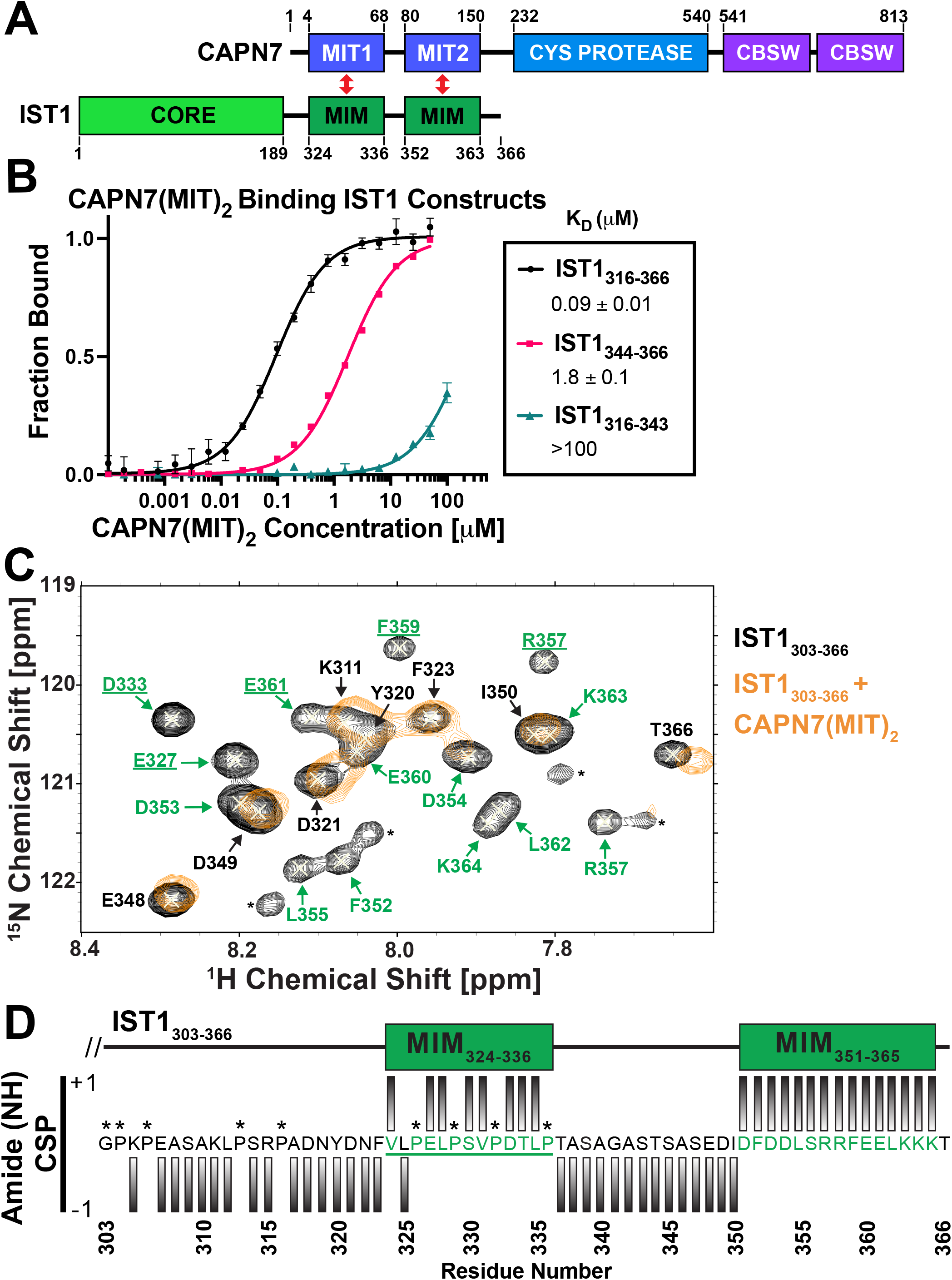
CAPN7 binds IST1 through tandem MIT domains. A) Domain organization of CAPN7 and IST1, depicting the binding interaction between tandem Microtubule Interacting and Trafficking (MIT) domains of CAPN7 and MIT Interacting Motif (MIM) elements of IST1 (red double headed arrows). Domain definitions: CORE, helical ESCRT-III core domain of IST1 which functions in filament formation and CBSW, tandem calpain-type beta sandwich domains of CAPN7. B) Fluorescence polarization anisotropy binding isotherms show CAPN7(MIT)_2_ binding to IST1 constructs spanning tandem or individual MIM elements (IST_316-366_, IST1_316-343_, and IST1_344-366_, respectively). Isotherm data points and dissociation constants are means from three independent experiments ± standard error. Error bars on the IST1_344-366_ isotherm are entirely masked by the data symbols. C) NMR chemical shift mapping of the CAPN7(MIT)_2_ binding sites on IST1_303-366_. Shown are a subset of overlaid HSQC spectra of free IST1_303-366_ (black contours) and IST1_303-366_ saturated with 1.3 molar equivalents of CAPN7(MIT)_2_ (orange contours). Amide NH resonances in the unbound state (black contours) without an accompanying bound state resonance (orange contours) identify those resonances with significant chemical shift perturbation upon CAPN7 binding (amino acid residue labels in green). In contrast, those resonances showing good superposition of unbound and bound (black and orange contours) identify those resonances with no chemical shift perturbation upon CAPN7 binding (amino acid residue labels in black) (Figure 1-figure supplement 1). D) Amide NH Chemical Shift Perturbations (CSP) mapped to IST1_303-366_ sequence. IST1 amide resonances are scored as “+1” for the first subset described above (23 amide resonances) and amino acid residues labeled with green or scored as “-1” (31) for the second subset described above (31 amide resonances) and amino acid residues labeled with black. Proline residues were not scored (asterisks) and G303 amide NH was not observable.

We used NMR chemical shift perturbation experiments to map IST1 binding residues that define a minimal IST1 peptide with full CAPN7(MIT)_2_ binding activity. NMR experiments employed a fully assigned ^15^N-labeled IST1_303-366_ peptide spanning both MIM elements that was titrated to saturation with unlabeled CAPN7(MIT)_2_. As expected, a subset of the IST1 amide (NH) resonances at the binding interface exhibited major perturbations upon titration; NH resonances decreased in intensity with increasing amounts of CAPN7(MIT)_2_ levels and were entirely absent upon saturation binding (Figure 1C and Figure 1 -figure supplement 1, compare black and orange peaks). In 23 cases, the NH resonances exhibited different chemical shifts in the bound vs. free complexes, exhibited slow exchange characteristics on the NMR time scale, and were significantly broadened in the bound complex (and therefore not visible). A second subset of 31 IST1 residues did not experience chemical shift environment changes and remained mobile upon CAPN7(MIT)_2_ binding (orange over black), implying that they did not contact CAPN7(MIT)_2_ in a stable manner. Residues in the former category mapped precisely to the two known IST1 MIM elements (Figure 1D), again implicating both elements in CAPN7(MIT)_2_ binding and implying that residues outside these elements likely do not contribute to binding. Consistent with this idea, a minimal IST1 construct (IST1_322-366_ K_D_ = 0.13± 0.01 μM) bound with the same affinity as the original construct (IST1_316-366_, K_D_ = 0.11 ± 0.01 μM) (Figure 1-figure supplement 2). In summary, our experiments demonstrate that both IST1 MIM elements contribute to CAPN7 binding and defined IST_322-366_ as a minimal CAPN7(MIT)_2_ binding construct used in subsequent structural studies.

### Crystal structure of CAPN7(MIT)_2_ – IST1 MIMs complex

We determined the crystal structure of the CAPN7(MIT)_2_ – IST1_322-366_ complex to define the interaction in molecular detail and to distinguish between the four possible binding modes in which individual CAPN7 MIT domains either bound simultaneously to both IST1 MIMs, as reported in other MIT:MIM interactions (Bajorek *et al*., 2009, Wenzel *et al*., 2022), or each CAPN7 MIT domain bound a different IST1 MIM element in one of two pairwise interactions. The structure revealed that each MIT domain binds a different MIM element, which is a configuration that has not been documented previously (Figure 2A and Table 1). The asymmetric unit contains two copies of the CAPN7(MIT)_2_–IST1_322-366_ complex, and the flexible linkers between the different MIT and MIM elements could, in principle, mediate two different possible connections between adjacent IST1 MIM elements (and their corresponding CAPN7 MIT domains). We have illustrated the simpler of the two models, which minimizes polypeptide chain cross-overs, but either connection is physically possible and the choice does not affect our interpretation of the model.

**Figure 2.**
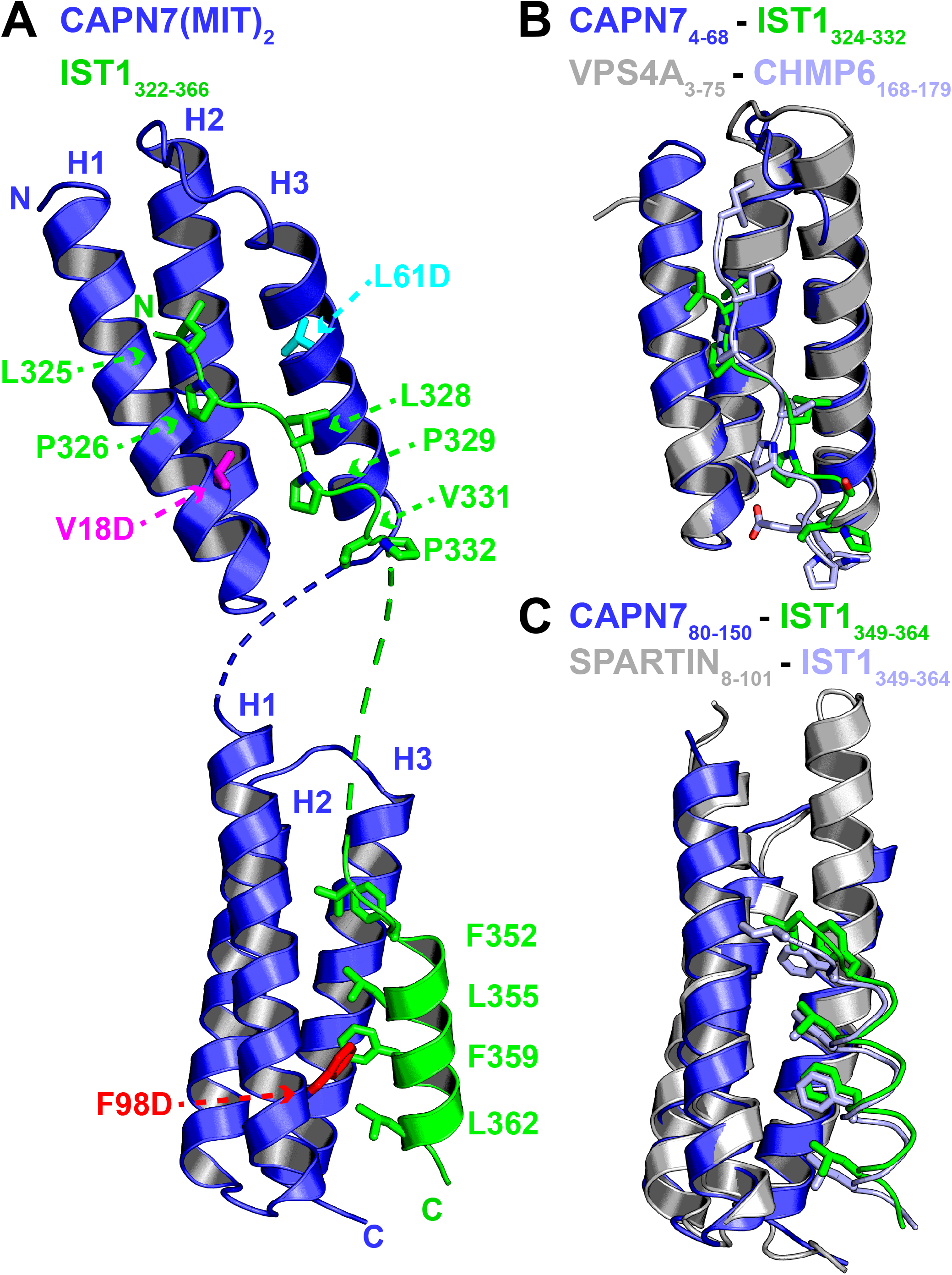
Crystal structure of the CAPN7(MIT)_2_-IST1_322-366_ complex. A) Ribbon representation of the CAPN7(MIT)_2_ (blue) in complex with IST1_322-366_ (green, with buried core interface sidechains shown) (PDB 8DFJ). Locations of residues that were mutated in CAPN7(MIT)_2_ are shown in magenta, turquoise, and red. Dashed lines show CAPN7 and IST1 residues that lack well defined electron density (see Materials and Methods). B) Structure of the CAPN7_4-68_–IST1_324-332_ complex (blue and green) overlaid with the structure of VPS4A_3-75_-CHMP6_168-179_ MIM complex (gray and light blue, PDB 2K3W). Note the similar binding Type 2 MIT-MIM binding modes. C) Structure of the CAPN7_80-150_-IST1_352-363_ complex (blue and green) overlaid with the structure of SPARTIN_8-101_-IST1_352-363_ complex (gray and light blue, PDB 4U7I) Note the similar Type 3 MIT-MIM binding modes.

**Table 1:**
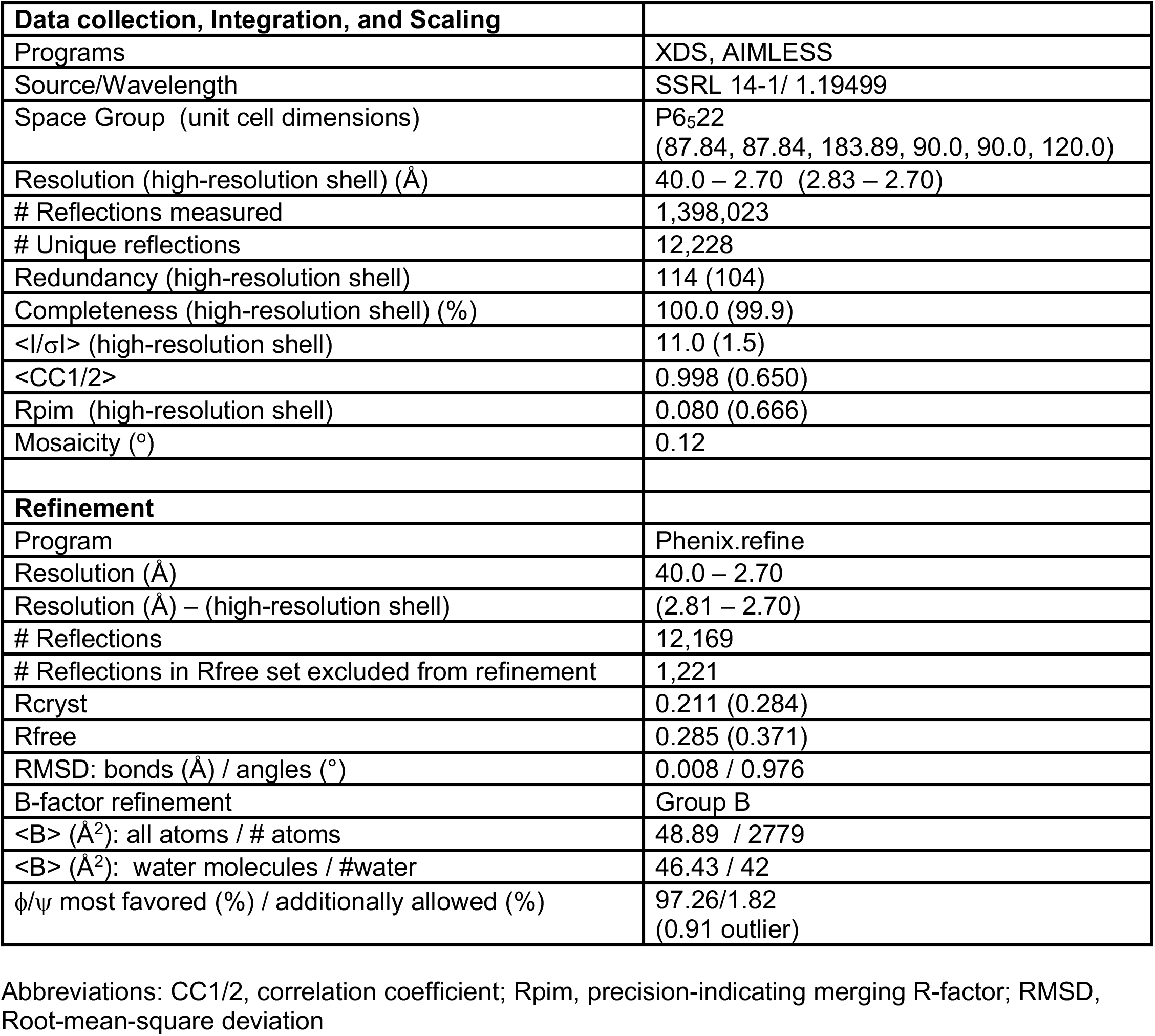
CAPN7(MIT)_2_ – IST1_322-366_ complex (PDB: 8DFJ) crystallographic data and refinement statistics.

Both CAPN7 MIT domains form the characteristic three-helix MIT bundle (Scott *et al*., 2005), and they are connected by a disordered six-residue linker. The N-terminal IST1 MIM element binds in an extended conformation in the helix 1/3 groove of the N-terminal CAPN7 MIT domain, making a canonical “Type-2” interaction (Figure 2A, top, and Figure 2-figure supplement 1A) (Kieffer *et al*., 2008; Samson *et al*., 2008; Vild and Xu, 2014; Kojima *et al*., 2016; Wenzel *et al*., 2022). The binding interface spans IST1 residues 325-332 (_325_VLPELPSVP_332_), which resembles the previously defined MIM2 consensus sequence (ΦPXΦPXXPΦP where Φ represents a hydrophobic residue and X a polar residue (Kojima *et al*., 2016)). Consistent with this observation, the overall structure closely resembles the Type-2 ESCRT-III-MIT complex formed by CHMP6_168-179_ bound to the VPS4A MIT domain (Figure 2B) (Kieffer *et al*., 2008; Skalicky *et al*., 2012).

Sequence divergence at either end of the MIM core appears to explain why the MIM2 element of CHMP6 binds preferentially to the VPS4A MIT domain, whereas the MIM2 element of IST1 prefers the first MIT domain of CAPN7. In the CAPN7-IST1 complex, IST1 Ser330 bulges away from the MIT domain to accommodate an intramolecular salt bridge formed by CAPN7 Asp21 and Arg63 (Figure 2-figure supplement 2A,C), and the peptide then bends back to allow Val331 and Pro332 to make hydrogen bonds and hydrophobic contacts, respectively. In the VPS4A MIT-CHMP6 complex, the CHMP6 Glu176 residue forms a salt bridge with VPS4A Lys23, whereas the equivalent interaction between CAPN7 MIT Gly24 and IST1 Val331 interaction has a very different character, thereby disfavoring CHMP6 binding to CAPN7 (Figure 2-figure supplement 2A,C). At the other end of the interface, helix 3 of the VPS4A MIT domain is two turns longer than the equivalent helix 3 in CAPN7 MIT (Figure 2B). This allows CHMP6 Ile168 to make a favorable interaction, whereas the shorter CAPN7 MIT helix 3 projects loop residue Leu50 directly toward the IST1_324-332_ N-terminus, in a position that would disfavor CHMP6 Ile168 binding (Figure 2-figure supplement 2A,B).

The C-terminal IST1 MIM element (_349_DIDFDDLSRRFEELKK_364_) forms an amphipathic helix that packs between helices 1 and 3 of the C-terminal CAPN7 MIT domain, making a canonical “Type-3” interaction (Figure 2A, bottom and Figure 2-figure supplement 1B) (Yang *et al*., 2008; Skalicky *et al*., 2012, Wenzel *et al*., 2022). Core residues from the hydrophobic face of the IST1 helix are buried in the CAPN7 MIT helix 1/3 groove, and the interface hydrophobic contacts, hydrogen bonds, and salt bridges are nearly identical to those seen when this same IST1 motif makes a Type-3 interaction with the MIT domain of SPARTIN (Figure 2C) (Guo and Xu, 2015, Wenzel *et al*., 2022). A previous study has reported the phosphorylation of Thr95 of CAPN7 (Mayya *et al*., 2009). This residue sits in the IST1_349-364_ binding site, adjacent to the detrimental F98D mutation (see below), and Thr95 phosphorylation would position the phosphate near Leu355 of IST1, creating an unfavorable electrostatic interaction and steric clash. We therefore anticipate that Thr95 phosphorylation would reduce IST1 binding and could negatively regulate the CAPN7-IST1 interaction.

The selectivity of each CAPN7 MIT domain for its cognate IST1 MIM element binding is dictated by the character of the MIT binding grooves. The hydrophobic contacts and hydrogen bonding potential of the residues within the N-terminal MIT domain are selective for Type-2 interactions (IST1_324-332_), whereas the C-terminal MIT domain selects for Type-3 interactions. This is consistent with our binding data for each individual CAPN7 MIT with each individual IST1 MIM element (Figure 2-figure supplement 3).

In summary, the CAPN7(MIT)_2_-IST1_322-366_ structure: 1) provides the first example of IST1_325-336_ bound to an MIT domain and confirms the expectation that this IST element engages in MIM2-type interactions, 2) demonstrates that IST1352-363 engages in a MIM3-type interaction when it binds CAPN7, and 3) shows that the separate IST1 MIM elements each bind distinct MIT domains within CAPN7(MIT)_2_.

### Mutational Analyses of the CAPN7 – IST1 complex

In our earlier study, we identified point mutations in the helix 1/3 grooves of each MIT domain that disrupt IST1 binding (Wenzel *et al*., 2022). The crystal structure of the CAPN7(MIT)_2_ – IST1_322-366_ complex explains why IST1 binding is disrupted by the V18D mutation in the first CAPN7 MIT domain (magenta, Figure 2A, top) and by the F98D mutation in the second MIT domain (red, Figure 2A, bottom). We also used the structure to design a control helix 1/2 mutation that was not expected to alter IST1 binding (L61D, cyan, Figure 2A, top). Fluorescence polarization anisotropy binding assays showed that the single V18D or F98D mutations in CAPN7(MIT)_2_ each reduced IST1_316-366_ binding >20-fold (Figure 3A), and that the V18D/F98D double mutation decreased IST1_316-366_ binding even further (>300-fold vs wt CAPN7(MIT)_2_). As predicted, the control L61D mutation did not affect IST1_316-366_ binding. Importantly, none of these mutations significantly disrupted the overall CAPN7(MIT)_2_ protein fold as assessed by circular dichroism spectroscopy (Figure 3-figure supplement 1A). Thus, we have identified CAPN7(MIT)_2_ mutations that diminish IST1316-366 complex formation by specifically disrupting each of the two MIT-MIM binding interfaces.

**Figure 3.**
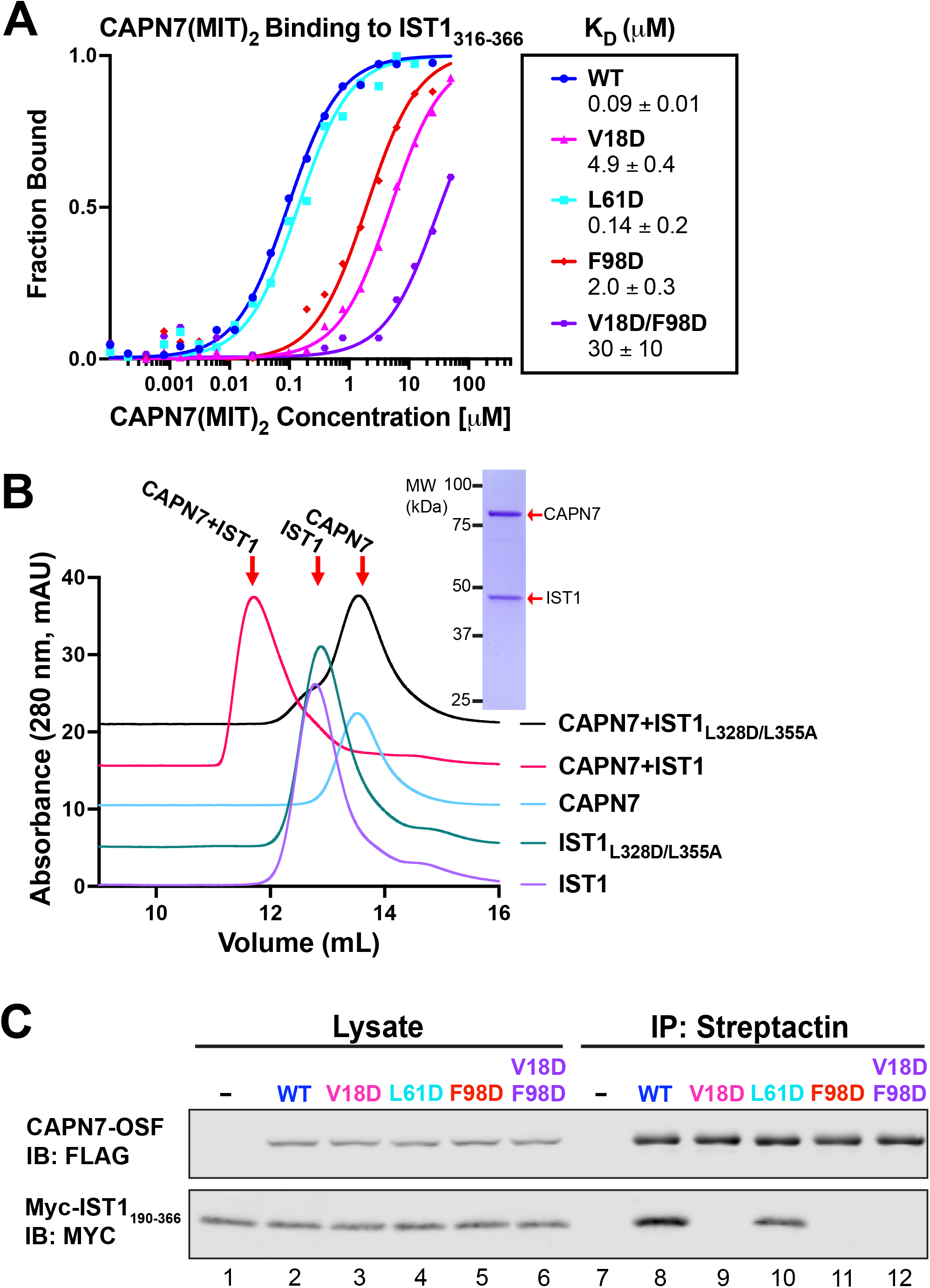
Mutational analyses of the CAPN7-IST1 complex. A) Fluorescence polarization anisotropy binding isotherms showing CAPN7(MIT)_2_ mutant constructs binding to IST1_316-366_. Isotherm data points and dissociation constants are means from three independent experiments ± standard error of mean. WT, V18D, and F98D binding isotherms are reproduced from Wenzel *et al*., 2022 for comparison. B) Size exclusion chromatography binding analysis of 1:1 mixtures of full-length CAPN7 binding to WT (black) or L328D/L355A mutant (green) full-length IST1. Note that the IST1 mutations disrupt CAPN7-IST1 complex formation. Elution profiles of each of the individual proteins are also shown for comparison. Inset image shows Coomassie stained SDS-PAGE gel of the peak fraction from the CAPN7+IST1 chromatogram. C) Co-immunoprecipitation of Myc-IST1_190-366_ with the indicated full-length CAPN7-OSF constructs from extracts of transfected HEK293T cells.

We also assessed the importance of these interfaces for association of full-length IST1 and CAPN7. The full-length proteins formed a 1:1 complex, as analyzed by size exclusion chromatography (Figure 3B). In contrast, inactivating mutations in the center of the two IST1 MIM elements interaction interfaces (L328D/L355A, see Figure 2A and Figure 3-figure supplement 2) blocked complex formation (Obita *et al*., 2007, Stuchell-Brereton *et al*., 2007, Kieffer *et al*., 2008, Bajorek *et al*., 2009), indicating that the crystallographically defined MIT-MIM interactions also mediate association of the full-length IST1 and CAPN7 proteins in solution. Finally, we analyzed the CAPN7-IST1 interaction in a cellular context using coimmunoprecipitation assays that employed our panel of different CAPN7 MIT mutations. Epitope-tagged CAPN7 and IST1 constructs were co-expressed in HEK293T cells, CAPN7-OSF constructs were immunoprecipitated from cellular extracts, and co-precipitated Myc-IST_190-366_ (Figure 3) or Myc-IST1 (Figure 3-figure supplement 1B) was detected by western blotting. Wildtype CAPN7 and the control CAPN7(L61D) mutant both co-precipitated IST1_190-366_ and IST1 efficiently, whereas all three inactivating CAPN7 MIT point mutations (V18D, F98D, and V18D/F98D) reduced the co-precipitation of Myc-IST_190-366_ or Myc-IST1 to control levels. Thus, the crystallographically defined MIT:MIM binding interfaces also mediate association of full length CAPN7 and IST1 in cells.

### IST1 recruits CAPN7 to the midbody

We next tested the midbody localization of wild-type CAPN7 and CAPN7 mutants lacking IST1 binding (V18D, F98D) or proteolytic (C290S) activities (Osako *et al*., 2010). These studies employed HeLa cell lines treated with siRNAs to deplete endogenous CAPN7, concomitantly induced to express integrated, siRNA-resistant, mCherry-tagged CAPN7 rescue constructs. Nup153 depletion was also used to maintain NoCut checkpoint signaling (Mackay *et al*., 2010; Strohacker *et al*., 2021; Wenzel *et al*., 2022), and thymidine treatment/washout was used to synchronize cell cycles and increase the proportion of midbody-stage cells (Figure 4-figure supplement 1). As expected, IST1 localized in a double-ring pattern on either side of the central Flemming body within the midbody (Agromayor *et al*., 2009; Bajorek *et al*., 2009), and wild-type CAPN7-mCherry colocalized with IST1 in the same pattern (Wenzel *et al*., 2022). CAPN7(C290S)-mCherry similarly colocalized with endogenous IST1 on either side of the Flemming body, whereas the two IST1 binding mutants did not (Figure 4A). IST1 midbody localization was normal in all cases. Quantification revealed that wild-type CAPN7 and CAPN7(C290S) colocalized with IST1 in ~80% of all IST1-positive midbodies, whereas the two IST1 non-binding mutants co-localized with IST1 in <5% of midbodies (Figure 4B) (Wenzel *et al*., 2022). Thus, IST1 recruits CAPN7 to midbodies, and efficient localization requires that both CAPN7 MIT domains bind both IST1 MIM elements, whereas CAPN7 proteolytic activity is not required for proper localization.

**Figure 4.**
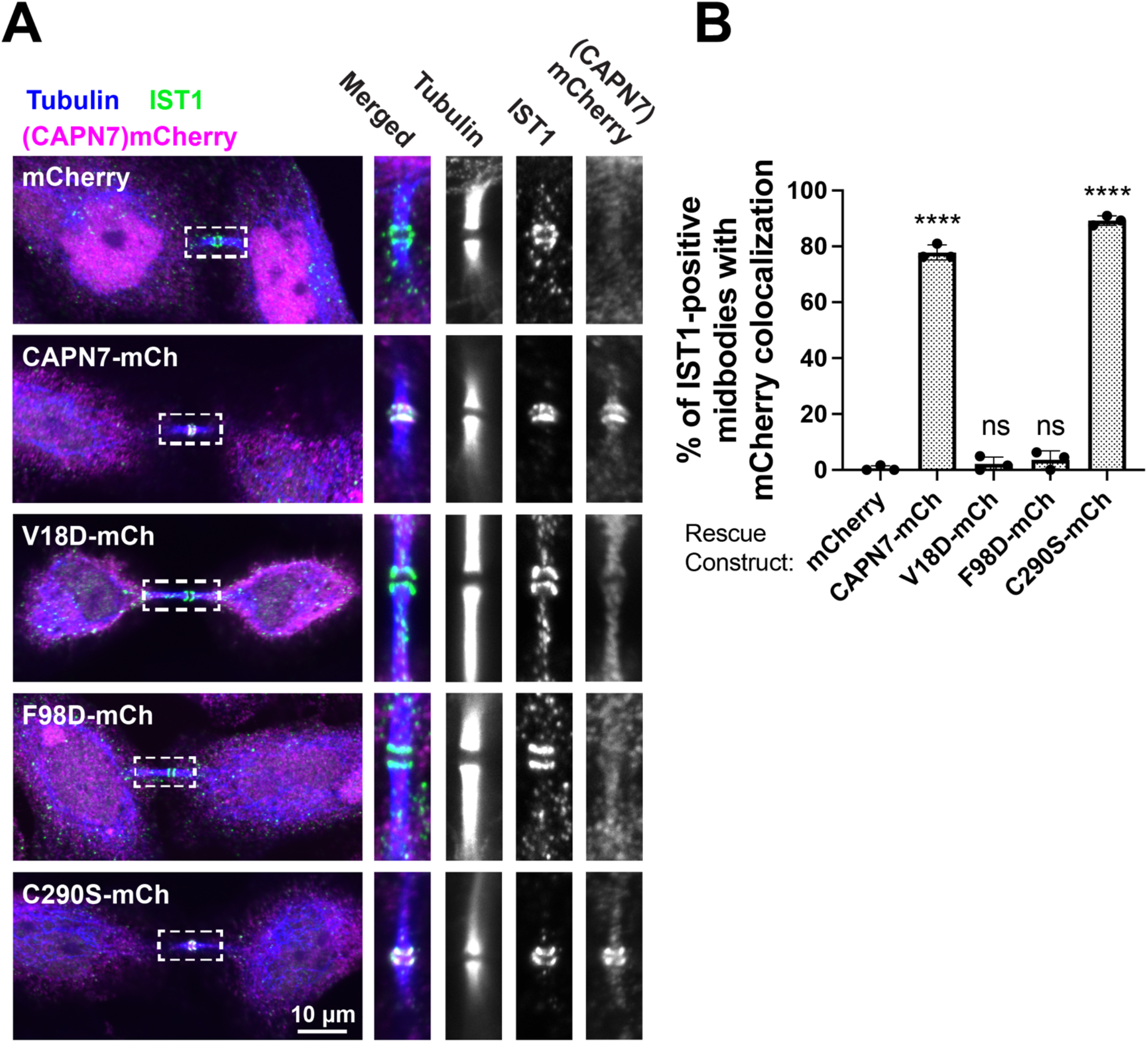
IST1 binding is required for CAPN7 midbody localization. A) Representative immunofluorescence images showing extent of midbody co-localization of mCherry-CAPN7 constructs (or the mCherry control), with endogenous IST1 in synchronous, NoCut checkpoint-activate cells. Endogenous CAPN7 was depleted with siRNA while inducibly expressing siRNA-resistant CAPN7-mCherry constructs. B) Quantification of the colocalization of mCherry-CAPN7 constructs with endogenous IST1 at midbodies (corresponding to the images in part A). Co-localization was scored blinded as described in the Materials and Methods. Bars represent the mean and standard error of mean from three independent experiments where >50 IST1-positive midbody-stage cells were counted per experiment. Statistical analyses were performed using unpaired t-tests that compared the percentage of rescue constructs that colocalized with IST1 at midbodies to the mCherry alone control. ****p<0.0001, ns (not significant) p>0.05.

### IST1-binding and proteolytic activity are required for CAPN7 functions in abscission and NoCut checkpoint maintenance

Our previous study indicated that CAPN7 promotes both abscission and NoCut checkpoint regulation (Wenzel *et al*., 2022). Here, we tested the requirements for CAPN7 to function in each of these processes. As expected, depletion of CAPN7 from asynchronous HeLa cells increased the fraction of cells that stalled or failed at abscission, as reflected by significant increases in midbody stage cells (from 5% to 10%) and multinucleate cells (from 6% to 22%) compared to control treatments (Figure 5A, and Figure 5-figure supplement 1A). Efficient abscission was almost completely rescued by expression of wild-type CAPN7-mCherry, whereas abscission was not rescued by CAPN7 constructs that were deficient in IST1-binding (V18D or F98D) or proteolysis (C290S). Thus, both IST1-binding and proteolytic activity are required for CAPN7 to support efficient abscission.

**Figure 5.**
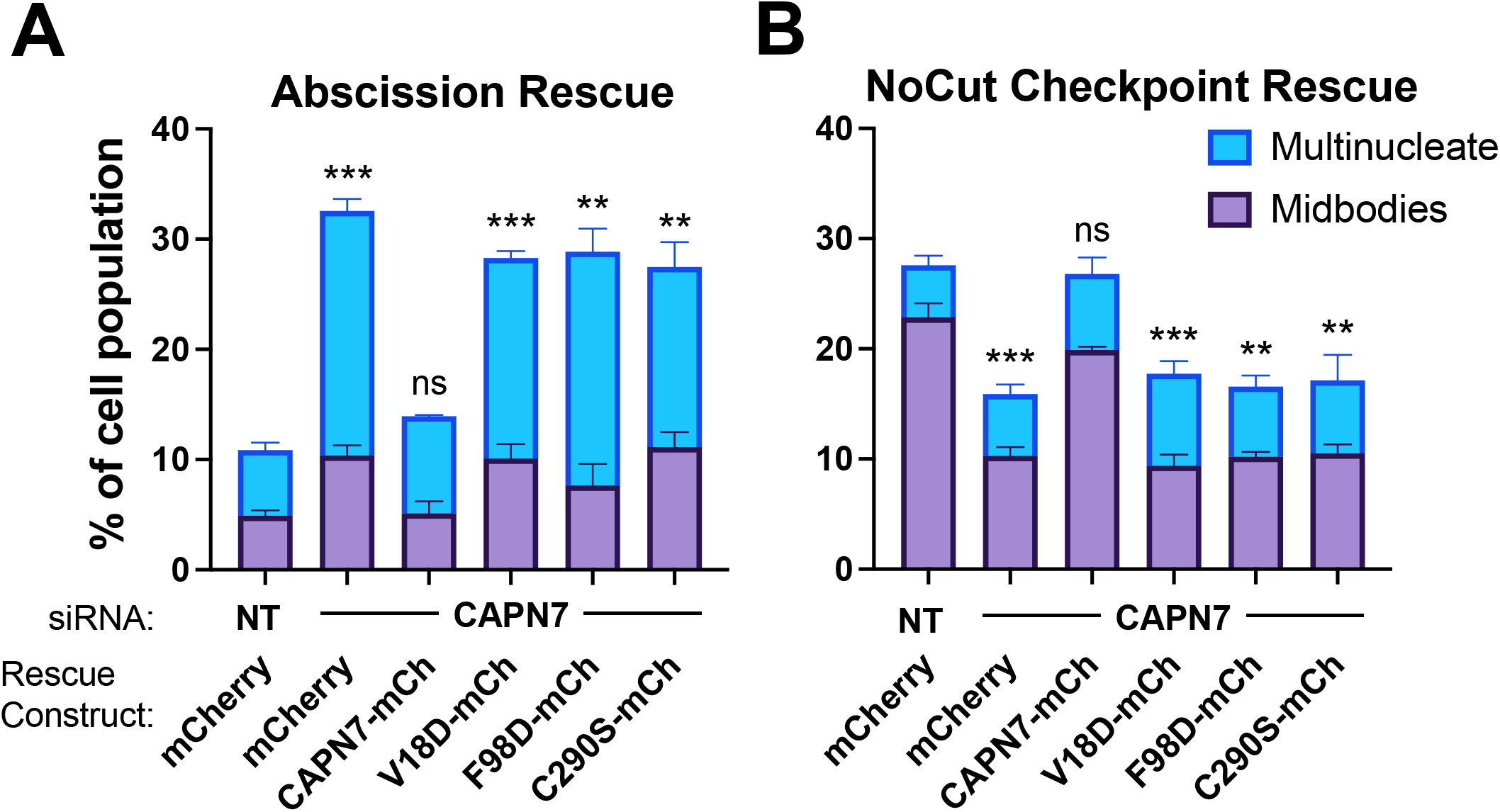
IST1-binding and catalytic activity are required for CAPN7 abscission and NoCut functions. (A, B) Quantification of midbody-stage and multinucleate HeLa cells from unperturbed asynchronous cultures (A), or cells in which NoCut checkpoint activity was sustained by Nup153 depletion (B). In all cases, cells were depleted of endogenous CAPN7, followed by expression of the designated DOX-inducible “rescue” construct. Bars represent the mean and standard error of mean from three independent experiments where >300 cells were counted per experiment. Statistical analyses were performed using unpaired t-tests comparing the sum of midbody-stage and multinucleate cells for each individual treatment to the same sum in siNT (non-targeting) control treated cells. ***p<0.001, **p<0.01, ns (not significant) p>0.05

We also tested the requirements for CAPN7 to support NoCut checkpoint signaling. In these experiments, NoCut signaling was maintained by Nup153 depletion, which delays abscission and raises the proportion of midbody-stage cells to 23% (Figure 5B and Figure 5-figure supplement 1B) (Mackay *et al*., 2010). Co-depletion of CAPN7 reduced the percentage of cells stalled at the midbody stage to 10%, indicating that CAPN7 is required to maintain the NoCut checkpoint signaling induced by Nup153 knockdown, in good agreement with our previous study (Wenzel *et al*., 2022). Rescue with wild-type CAPN7-mCherry restored NoCut checkpoint function almost completely (20% midbodies), whereas the IST1-binding or catalytically-dead mutant CAPN7 constructs failed to rescue the checkpoint (all ~10% midbodies). Hence, CAPN7 must bind IST1 and be an active protease to participate in NoCut checkpoint regulation.

The dual requirements for CAPN7 proteolysis in the competing processes of abscission and NoCut abscission delay are counterintuitive, but we envision that CAPN7 could participate in both processes if substrate specificities or activities change with the different checkpoint statuses of the cell; for example, through altered post-translational modifications or binding partners.

### CAPN7 as an evolutionarily conserved ESCRT-signaling protein

Our experiments imply that proteolysis is critical for different CAPN7 functions, but physiological substrates have not yet been identified. GFP-CAPN7 can perform autolysis, and this activity is enhanced by IST1 binding (Osako *et al*., 2010), implying that interactions with IST1 at the midbody could activate CAPN7 proteolytic activity.

In addition to our studies implicating CAPN7 midbody functions during cytokinesis, other biological functions for CAPN7 have also been proposed, including in ESCRT-dependent degradation of EGFR through the endosomal/MVB pathway (Yorikawa *et al*., 2008; Maemoto *et al*., 2014), and in human endometrial stromal cell migration and invasion (Liu *et al*., 2013) and decidualization (Kang *et al*., 2022). Aberrant CAPN7 expression may also impair embryo implantation through a currently unknown mechanism (Yan *et al*., 2018).

CAPN7 has evolutionarily conserved orthologs throughout Eukaryota, including in Aspergillus and yeast, where its role in ESCRT-dependent pH signaling is well described (Denison *et al*., 1995; Orejas *et al*., 1995; Futai *et al*., 1999; Peñalva *et al*., 2014). In Aspergillus, pH signaling requires proteolytic activation of the transcription factor PacC (no known human ortholog), which initiates an adaptive transcriptional response. PacC is cleaved and activated by the MIT domain-containing, Calpain-like CAPN7 ortholog protease PalB, which is recruited to membrane-associated ESCRT-III filaments in response to alkaline pH through a MIT:MIM interaction with the ESCRT-III protein Vps24 (CHMP3 ortholog) (Rodríguez-Galán *et al*., 2009). These ESCRT assemblies also contain PalA (ALIX ortholog), which co-recruits PacC to serve as a PalB substrate (Xu and Mitchell, 2001; Vincent *et al*., 2003).

The pH signaling system in fungi has striking parallels with our model for human CAPN7 functions in cytokinesis, including: 1) MIT:MIM interactions between CAPN7 and ESCRT-III, which serve to recruit the protease to a key ESCRT signaling site. 2) The local presence of the homologous PalA and ALIX proteins. In the pH sensing system, PalA is thought to act as a scaffold that “presents” the substrate to PalB for proteolysis on ESCRT-III assemblies, thus providing an extra level of spatiotemporal specificity for proteolysis. In the mammalian midbody, ALIX nucleates ESCRT-III filament formation. 3) A requirement for proteolysis in downstream signal transduction. Thus, ESCRT-III assemblies are apparently evolutionarily conserved platforms that recruit and unite active calpain proteases and their substrates with spatiotemporal specificity to regulate key biological processes.

## Materials and Methods

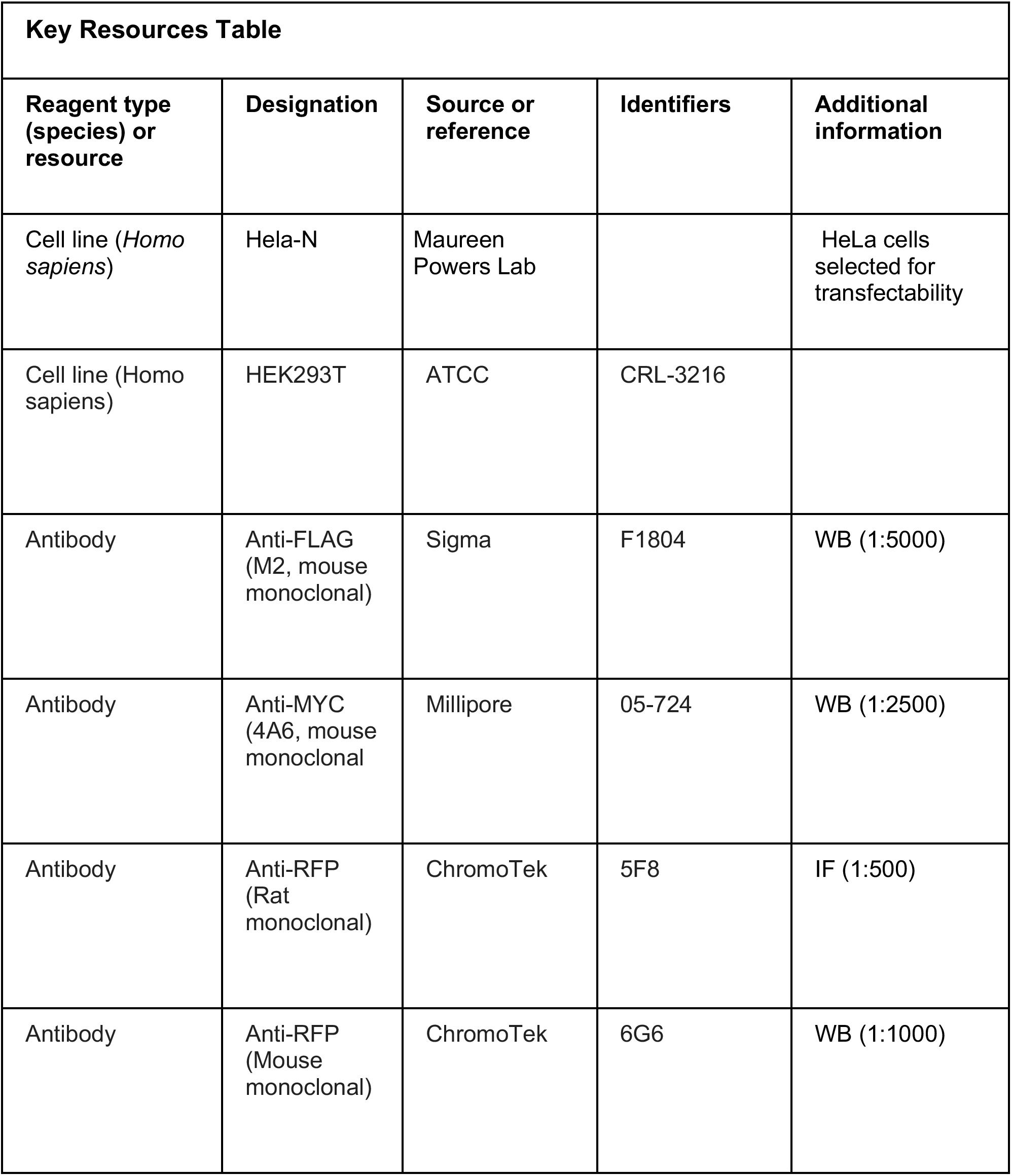
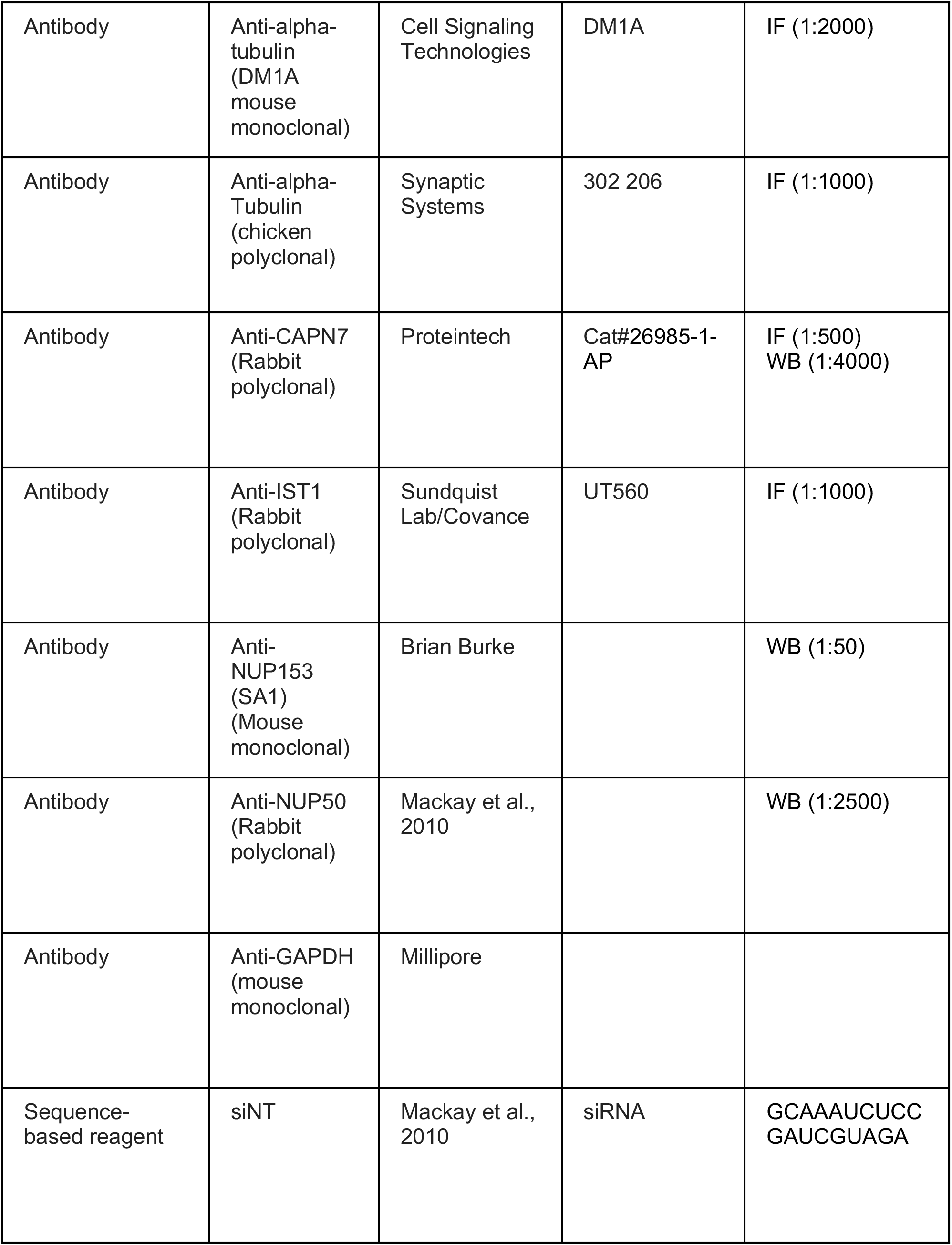
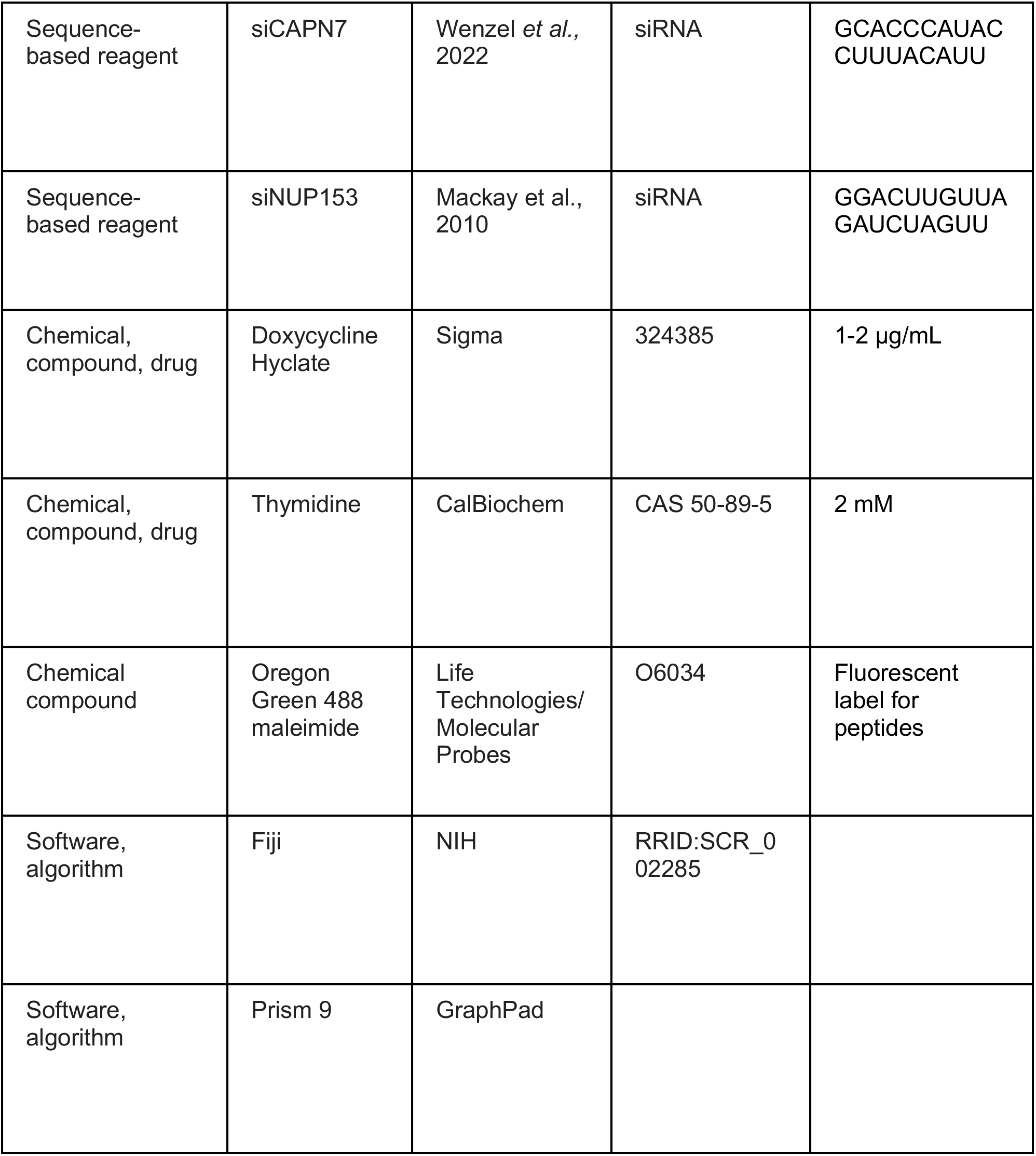

### Cloning

All plasmids created for this study were made by Gibson Assembly (Gibson *et al*., 2009) using vectors linearized by restriction enzymes (New England Biolabs). pCA528 bacterial expression plasmids expressing His_6_-SUMO-tagged fusion proteins with a native N-terminus after tag cleavage were linearized with BsaI and BamHI. pCAG-OSF mammalian expression plasmids used in co-IP binding assays were linearized with KpnI and XhoI. pLVX vectors used for generating inducible cell lines were linearized by BamHI and MluI. Point mutations were generated using QuikChange II Site-directed mutagenesis kit (Agilent). See Supplementary Table 2 for complete plasmid list details.

### Bacterial expression of (His)_6_-SUMO-fusion proteins

Proteins were expressed in BL21 RIPL cells grown in ZYP-5052 autoinduction media (Studier, 2005). Transformed cells were initially grown for 3-6 h at 37 °C to an OD600 of 0-4-0.6, and then switched to 19 °C for an additional 20 h. Cells were harvested by centrifugation at 6,000 x g and cell pellets were stored at −80°C.

### Purification of (His)_6_-SUMO-fusion proteins

All purification steps were carried out at 4°C except where noted. Frozen cell pellets were thawed and resuspended in lysis buffer: 50 mM Tris (pH 8.0 at 23 °C), 500 mM NaCl, 2 mM Imidazole, 1 mM Dithioreitol (DTT), supplemented with 0.125% sodium deoxycholate, lysozyme (25 μg/mL), PMSF (100 μM), Pepstatin (10 μM), Leupeptin (100 μM), Aprotinin (1 μM), DNAseI (10 μg/mL) and 2 mM MgCl_2_. Cells were lysed by sonication and lysates were clarified by centrifugation at 40,000 x g for 45 min. Clarified supernatant was filtered through a 0.45 μM PES syringe filter and incubated with 10 mL of cOmplete His-Tag purification beads (Roche) for 1 h with gentle rocking. Beads were washed with 500 mL wash buffer: 50 mM Tris (pH 8.0 at 23 °C), 500 mM NaCl, 2 mM Imidazole, 1 mM DTT. Fusion proteins were eluted with 50 mL of wash buffer supplemented with 250 mM Imidazole. Eluted proteins were treated with 100 ug His_6_-ULP1 protease overnight in 6-8 k MWCO dialysis bags while dialyzing against 2 x 2 L of 25 mM Tris pH (8.0 at 23 °C), 100 mM NaCl, 1 mM TCEP, 0.5 mM EDTA. The dialysate was purified by Q Sepharose chromatography (HiTrap Q HP 5 mL; Cytiva Life Sciences) with a linear gradient elution from 100-1000 mM NaCl. Fractions containing cleaved fusion protein were then passed over 5 mL of cOmplete His-Tag purification beads to remove residual His_6_-Sumo-fusion protein and His_6_-Sumo cleaved tag. Proteins were concentrated and purified by Superdex 75 or Superdex 200 gel filtration chromatography (120 mL;16/600; Cytiva Life Sciences) in 25 mM Tris (pH 7.2 at 23 °C), 150 mM NaCl, 1 mM TCEP, and 0.5 mM EDTA. Highly pure fractions were pooled and concentrated. Masses of each protein was confirmed with ESI-MS (University of Utah Mass Spectrometry Core Facility, see Supplemental Table 1). Yields ranged from 0.2-50 mg/L of bacterial culture. Full-length CAPN7 yields were poor (~0.2 mg/L), hence full-length IST1 and its mutants (~20 mg/L) were used for gel filtration binding experiments in Figure 3B.

### Peptide fluorescent labeling

Fluorescent labeling was performed by the University of Utah DNA/Peptide Synthesis Core as described previously (Caballe et al., 2015; Talledge et al., 2018). Briefly, peptides were labeled in DMSO using ~1.3-fold molar excess of Oregon Green 488 maleimide (Life Technologies/Molecular Probes #O6034) dissolved in a 1:1 solution of acetonitrile:DMSO. Reverse phase HPLC was used to monitor the reactions and separate labeled peptides from unreacted dye and unlabeled peptide. Labeled peptide fractions were dried under vacuum and dissolved in water. Peptide concentrations were quantified using the absorbance of Oregon Green 488 at 491 nm (*ϵ* = 83,000 cm^-1^ M^-1^ in 50 mM potassium phosphate, pH 9.0).

### Fluorescence polarization anisotropy binding assays

Binding experiments were performed as described previously (Caballe et al., 2015) in 25 mM Tris, pH 7.2, 150 mM NaCl, 0.1 mg/mL Bovine Serum Albumin (BSA), 0.01% Tween-20, and 1 mM DTT, with 250 pM fluor-labeled IST1 peptides and purified CAPN7 MIT domain constructs at the indicated concentrations. A Biotek Synergy Neo Multi-Mode plate reader was used to measure fluorescence polarization with excitation at 485 nm and emission (detection) at 535 nm. Binding isotherms were fit to 1:1 models using Prism 9 (GraphPad). Reported K_D_ values are averages from at least three independent isotherms ± standard error mean. No attempt was made to estimate K_D_ if a binding isotherm did not reach half bound at the highest concentration used and in these cases the K_D_ is listed as greater than the highest concentration used. The K_D_ is listed as ND (Not Determined) if the change in fluorescence polarization was too small for curve fitting (as in Figure 2 – figure supplement 3).

### NMR Spectroscopy

NMR data were collected and processed as described previously for ^15^N-enriched IST1_303-366_ (Caballe *et al*., 2015). Chemical shift perturbations induced by CAPN7 binding to IST1 were identified by titrating 0.25 mM of uniformly ^15^N-labeled IST1(303-366) with unlabeled CAPN7(MIT)_2_ at molar equivalents of 0.25, 0.5, 1.0, and 1.3. IST1 amide NH resonances that decreased in intensity with increasing levels of CAPN7(MIT)_2_ were scored as “+1” (23 aa). Resonances with small or no shifts (< ½ linewidth at half height) and similar resonance intensity were scored as “-1” (31 aa). Prolines are absent amide NH resonances and Gly303 is not observable; both are scored as “0” (9 aa).

Amide NH resonances scored as “+1” are in NMR slow exchange, experience different magnetic environments in free vs. bound states, are broadened beyond detection in the saturated bound complex, and define those IST1 residues at binding sites. Resonances scored as “-1” are in NMR fast exchange, display small or no chemical shift perturbations, remain flexible in free and bound complexes, and define nonbinding regions of IST1.

### X-Ray crystallography

In preparation for crystallization, purified CAPN7_1-165_ (~60 mg/ml) and IST1_322-366_ (~20 mg/mL), in a buffer containing 25 mM Tris (pH 7.2) at 23 °C, 150 mM NaCl, 1 mM TCEP, 0.5 mM EDTA, were mixed in a 1:2 molar ratio (CAPN7:IST1) so that the final concentration of CAPN7_1-165_ was 20 mg/mL. Crystallization was carried out in vapor-diffusion crystallization trays with the aid of an ARI Gryphon liquid-handling robot (Art Robbins Instruments). Flat hexagonal plates measuring about 100-200 μm across grew after about 3 weeks at 21 °C in the Rigaku Wizard Cryo screen, condition D5 (25% (v/v) 1,2-Propanediol, 100 mM Sodium phosphate dibasic/ Citric acid pH 4.2, 5% (w/v) PEG 3000, 10% (v/v) Glycerol). In preparation for data collection, crystals were transferred briefly (less than 20 seconds) into mother liquor with 25% added glycerol, suspended in a small nylon sample mounting loop, and cryocooled by plunging into liquid nitrogen.

X-ray diffraction data were collected at the Stanford Synchrotron Radiation Lightsource (SSRL). During data collection the crystal was maintained at 100 K with the aid a cold nitrogen gas stream. Data were integrated and scaled using XDS (Kabsch, 2010a; Kabsch 2010b) and AIMLESS (Evans, 2011; Evans and Murshudov, 2013) (Table 1). Initial phases were obtained using phaser in the PHENIX software suite (Bunkoczi *et al*., 2013) using VPS4B MIT (PDB 4U7Y) (Guo and Xu, 2015) as a search model. The resulting electron density maps were readily interpretable, allowing a model to be built using Coot (Emsley and Cowtan, 2004; Emsley *et al*., 2010), and refined with phenix.refine (Liebschner *et al*., 2019).

Model validation of the CAPN7 MIT domains was performed by generating an omit map (FoFc) demonstrating unbiased density for Phe and Tyr residues where these side chains were omitted from the model prior to refinement and calculation of model-based phases. Model validation of the IST1 MIM elements was performed by omitting all IST1 residues (Figure 2-Figure supplement 1). The final model was refined against all data to R_work_=0.211 and R_free_=0.285. Full refinement statistics and details can be found in Table 1. Structure coordinates have been deposited in the RCSB Protein Data Bank (PDB 8DFJ).

Structure alignments shown in Figure 2 were generated using lsqkab (Kabsch, 1976) in the CCP4 program suite (Winn *et al*., 2011).

Protein interfaces and details of protein-protein contacts were analysed with PISA (Krissinel and Henrick, 2007) and LigPlot+ (Laskowski and Swindells, 2011).

### Cell culture

HEK293T and HeLa cells were cultured at 37 °C and 5% CO_2_ in high glucose DMEM (Gibco) supplemented with 10% FBS. TetOn-HeLa cells were additionally supplemented with 500 μg/mL G418 (Invitrogen) to maintain expression of the Tet-On Advanced protein. Doxycycline-inducible CAPN7 expression construct cell lines were generated in the parental TetOn-Advanced cell line and further supplemented with 0.5 μg/mL puromycin (Invitrogen).

### Cell lines

Cell lines were authenticated, generated, and validated as described previously (Wenzel *et al*., 2022). Briefly, the parental HeLa cell lines were authenticated by genomic sequencing of 24 loci (University of Utah Sequencing Core) and confirmed to be mycoplasma-free by routine PCR testing (ABM) following the manufacturer’s protocol. To generate stable cell lines with doxycycline-inducible expression, the parental HeLa Tet-On Advanced cells were transfected with pLVX-tight puro plasmids containing the CAPN7 constructs (see Supplemental Table 2) and selected for 14 days in 500 μg/mL G418 + 1 μg/mL puromycin. Single colonies were expanded and screened for expression by immunofluorescence and western blotting. Selected clones were further validated by sequencing the PCR amplified exogenous CAPN7 construct. Protein expression was induced by addition of 1 – 2 μg/mL doxycycline.

### Gel filtration chromatography binding assay

Highly pure (>99%) individual proteins were resolved by gel filtration chromatography at 4 °C by injecting 2 nmol protein in a 100 uL sample loop onto a Superdex 200 (24 mL; 10/300 GL, Cytiva Life Sciences) at 0.5 mL/min flow rate using 20 mM Tris (pH 7.2 at 23 °C), 150 mM NaCl, 0.5 mM TCEP as eluent and binding buffer. Protein elution was monitored by UV absorbance at 280 nm. The column was calibrated using molecular weight standards (BioRad).

Protein complexes were made by combining the two components in a 1:1 molar ratio to a final concentration of 20 μM in binding buffer and incubated on ice for 1 h. Protein complexes were spun down at 16,000 x g for 10 min at 4 °C, then 2 nmol were immediately injected onto the column. Elution fractions were collected and the fraction corresponding to the peak of CAPN7-IST1 chromatogram was run on a Coomassie-stained SDS-PAGE gel to produce the inset image in Figure 3B.

### Co-immunoprecipitation experiments

HEK293T cells were seeded at 0.5 x 10^6^ cells per well in 6-well plates and transfected about 24 h later when cell confluency was ~80% with 3 μg DNA complexed with PEI. DNA mixtures contained 1.5 μg of plasmids encoding either pCAG-Myc-IST1(190-366) or pCAG-Myc-IST1(1-366) plus 1.5 μg of plasmids encoding either empty vector (pCAG-MCS2-OSF) or the pCAG-CAPN7 constructs (See Supplementary Table 2). Cells were harvested 48 h post transfection and lysed in 500 μL cold lysis buffer (50 mM Tris (pH 7.2 at 23 °C), 150 mM NaCl, 0.5% TritonX-100, 1 mM DTT) supplemented with mammalian protease inhibitor cocktail (1:100, Sigma). Cells were lysed for 15 minutes on ice with brief vortexing every 5 minutes. Lysates were clarified by centrifugation at 16,000 x g for 10 min at 4 °C, then applied to 20 μL of a 50% slurry of Streptactin resin (IBA Biosciences) for 1 h at 4 °C. Beads were washed three times with 500 μL lysis buffer and aspirated to near dryness. Bound proteins were eluted by boiling the Streptactin resin in 40 μL of 2x Laemmli sample buffer, resolved by SDS-PAGE, and detected by Western blotting.

### siRNA transfections

For experiments in Figure 4, transfection protocols were as follows: Day 1 – 0.5 x 10^5^ cells were seeded on fibronectin-coated coverslips in a 24-well plate for immunofluorescence or 3 x 10^5^ cells in a 6-well plate for western blotting and transfected with 20 nM siCAPN7 and 10 nM siNup153 complexed with Lipofectamine RNAiMAX (Invitrogen) (See Supplementary Table 3 for sequences) with 2 μg/mL doxycycline. Eight hours later, the media was changed with 2 μg/mL doxycycline and 2 mM thymidine. Day 2 – 24 hours later, the thymidine was washed out with three washes of PBS and fresh media was added with 2 μg/mL doxycycline. Day 3 – Cells were harvested 16 h after thymidine washout for immunofluorescence and immunoblotting.

For experiments in Figure 5A, transfection protocols were performed with 72 h siRNA knockdown and 48 h doxycycline-induced CAPN7 construct expression: Day 1 – 5 x 10^5^ cells were seeded in a 6-well plate and transfected with 20 nM siNT or siCAPN7 complexed with RNAiMAX (see Supplementary Table 3 for sequences). Day 2 – 24 h later, cells were split onto either fibronectin-coated coverslips (0.5 x 10^5^ cells) for immunofluorescence or a 6-well plate for immunoblotting (2.5 x 10^5^ cells) and transfected again with 20 nM siNT or siCAPN7 with 1 μg/mL doxycycline. Day 3 – 24 h later, media was changed with 1 μg/mL doxycycline. Day 4 – 24 h later, cells were harvested for immunofluorescence and immunoblotting.

For experiments in Figure 5B, transfection protocols were performed with 72 h siRNA knockdown of CAPN7 and 48 h knockdown of Nup153 to induce a checkpoint arrest and 72 h doxycycline-induced CAPN7 construct expression: Day 1 – 5 x 10^5^ cells were seeded in a 6-well plate and transfected with 20 nM siNT or siCAPN7 complexed with RNAiMAX with 2 μg/mL doxycycline. Day 2 – 24 h later, cells were split onto coverslips or 6-well plate as above with 20 nM siNT or siCAPN7 plus 10 nM siNT or siNup153 with 2 μg/mL doxycycline. Day 3 – 24 h later, media was changed with 2 μg/mL doxycycline. Day 4 – 24 h later, cells were harvested for immunofluorescence and immunoblotting.

### Immunoblotting

Immunoblots related to Figures 4 and 5 were performed as previously described (Wenzel *et al*., 2022). Briefly, Cells were lysed in RIPA buffer (Thermo Fisher) supplemented with mammalian protease inhibitor cocktail (1:100, Sigma) for 15 minutes on ice with brief vortexing every 5 minutes. Lysates were cleared by centrifugation at 17,000 x g for 10 minutes at 4 °C. Lysate protein concentrations were determined using the BCA Assay (Thermo Fisher) before SDS-PAGE. 10 μg lysate per sample were prepared with SDS loading buffer, resolved by SDS-PAGE, and transferred to PVDF membranes. Membranes were blocked for 1 hour at room temperature in 5% milk in TBS, then incubated overnight at 4 °C with primary antibodies (see Supplemental Table 4 for dilutions). Following 3 x 10-minute washes in TBS-T, membranes were incubated with the corresponding secondary antibodies for 1 hour at 23 °C, washed again with TBS-T, and imaged using a LiCor Odyssey infrared scanner.

### Immunofluorescence imaging and phenotype quantification

Cells were seeded on fibronectin-coated glass coverslips in 24-well plates and treated with siRNAs as described above. Coverslips were harvested by washing once with PBS and once with 1X PHEM (25 mM HEPES, 60 mM PIPES, 10 mM EGTA, 4 mM MgCl_2_, pH 6.9), then fixed and permeabilized in 4% PFA, 1X PHEM, 0.5% TritonX-100 for 15 min at 23 °C. Coverslips were then washed 3x with PBS-T (0.1% TritonX-100) and blocked for 1 h in 5% normal donkey serum in PBS-T. Coverslips were then incubated with primary antibody for 1 – 2 h at 23 °C, diluted in 1% BSA in PBS-T (see Supplementary Table 4 for antibody dilutions). Coverslips were then washed 3 x PBS-T, then incubated with secondary antibodies for 1 h at 23 °C, diluted in 1% BSA in PBS-T (See Supplementary Table 4 for antibody dilutions). Coverslips were washed 3 x PBS-T, once with water, then mounted in ProLong Diamond (Invitrogen) for localization experiments in Figure 4 or ProLong Diamond with DAPI for functional experiments in Figure 5. Mounted coverslips were cured for at least 24 h before imaging.

Images for Figure 4A were acquired on a Leica SP8 white light laser confocal microscope using a 63x 1.4 oil HC PL APO objective. Images were acquired as Z-stacks and presented as maximum intensity Z-projections using the Leica App Suite X Software.

Images for Figure 4B and 5A,B phenotype quantification were acquired as previously described (Wenzel *et al*., 2022). Briefly, images were acquired using a Nikon Ti-E inverted microscope equipped with a 60x PlanApo oil immersion objective, an Andor Zyla CMOS camera, and an automated Prior II motorized stage controlled with the Nikon Elements software. For phenotype quantification in Figure 4 and 5, the software was used to acquire 25 images using a randomized 5×5 grid pattern. The images were then blinded and scored to reduce any potential for bias. For Figure 4B localization quantification, only midbodies with in-focus IST1 staining were scored for the obvious presence or absence of mCherry signal. Quantification for Figures 4B, 5A, and 5B were performed from three independent, biological replicates (cells seeded and treated on different days). Quantification and statistical analyses were performed using GraphPad Prism 9.

### Materials and Data Availability

X-Ray diffraction data were deposited in the PDB under accession code 8DFJ. NMR chemical shift assignments for IST1_303-366_ are available at Biological Magnetic Resonance Bank (Accession no: 25393)(Caballe *et al*., 2015). All new plasmids generated for this study have been deposited at Addgene.

## Acknowledgements

We thank Courtney Dailley for assistance purifying proteins and performing gel filtration chromatography. Oligonucleotides and peptides (IST1_316-343_ and IST1_344-366_) were synthesized and purified by HPLC by the DNA/Peptide Facility, part of the Health Sciences Center Cores at the University of Utah and we thank Mike Hanson for assistance. DNA sequencing was performed by the DNA sequencing core facility, University of Utah and we thank Derek Warner and Michael Powers for assistance. We thank the Cell Imaging Core at the University of Utah for use of equipment (Nikon Ti-E and Leica SP8 microscopes) and thank Xiang Wang for his assistance. Proteomics mass spectrometry analysis was performed at the Mass Spectrometry and Proteomics Core Facility at the University of Utah and we thank Sandra Osburn for assistance. Mass spectrometry equipment was obtained through a Shared Instrumentation Grant 1 S10 OD018210 01A1. This work was funded by grants from NIH 5R01GM112080 (WIS, KSU, CPH) and F31GM139318 (ELP). Use of the Stanford Synchrotron Radiation Lightsource, SLAC National Accelerator Laboratory, is supported by the U.S. Department of Energy, Offices of Science, Office of Basic Energy Sciences under Contract No. DE-AC02-76SF00515. The SSRL Structural Molecular Biology Program is supported by the DOE Office of Biological and Environmental Research, and by the NIH, National Institute of General Medicine Sciences (including P41GM103393). The contents of this publication are solely the responsibility of the authors and do not necessarily represent the official views of NIGMS or NIH.

---

**Supplemental Table 1.** ESI-MS mass confirmation of purified proteins.

| Expression plasmid | Expected Mass (Da) | Measured Mass (Da) | Uniprot # | Notes |
| --- | --- | --- | --- | --- |
| pCA528 CAPN7 (1-165) | 18579.80 | 18581.34 | Q9Y6W3 | Wenzel *et al.*, 2022 |
| pCA528 CAPN7 V18D (1-165) | 18595.75 | 18595.35 | Q9Y6W3 | Wenzel *et al.*, 2022 |
| pCA528 CAPN7 L61D (1-165) | 18581.73 | 18581.35 | Q9Y6W3 |  |
| pCA528 CAPN7 F98D (1-165) | 18547.41 | 18547.35 | Q9Y6W3 | Wenzel *et al.*, 2022 |
| pCA528 CAPN7 V18D,F98D (1-165) | 18562.31 | 18562.37 | Q9Y6W3 |  |
| pCA528 CAPN7 (1-813) | 92652.41 | 92650.89 | Q9Y6W3 |  |
| pCA528 IST1 (316-366) (C-Cys) | 6252.69 | 6251.69 | P53390-4 | Non-native N-terminal Gly, reacted with Oregon Green. Wenzel *et al.*, 2022 |
| pCA528 IST1 (322-366) | 4895.46 | 4895.45 | P53390-4 |  |
| pCA528 IST1 (1-366) | 39978.83 | 39978.54 | P53390-4 |  |

**Supplemental Table 2.**
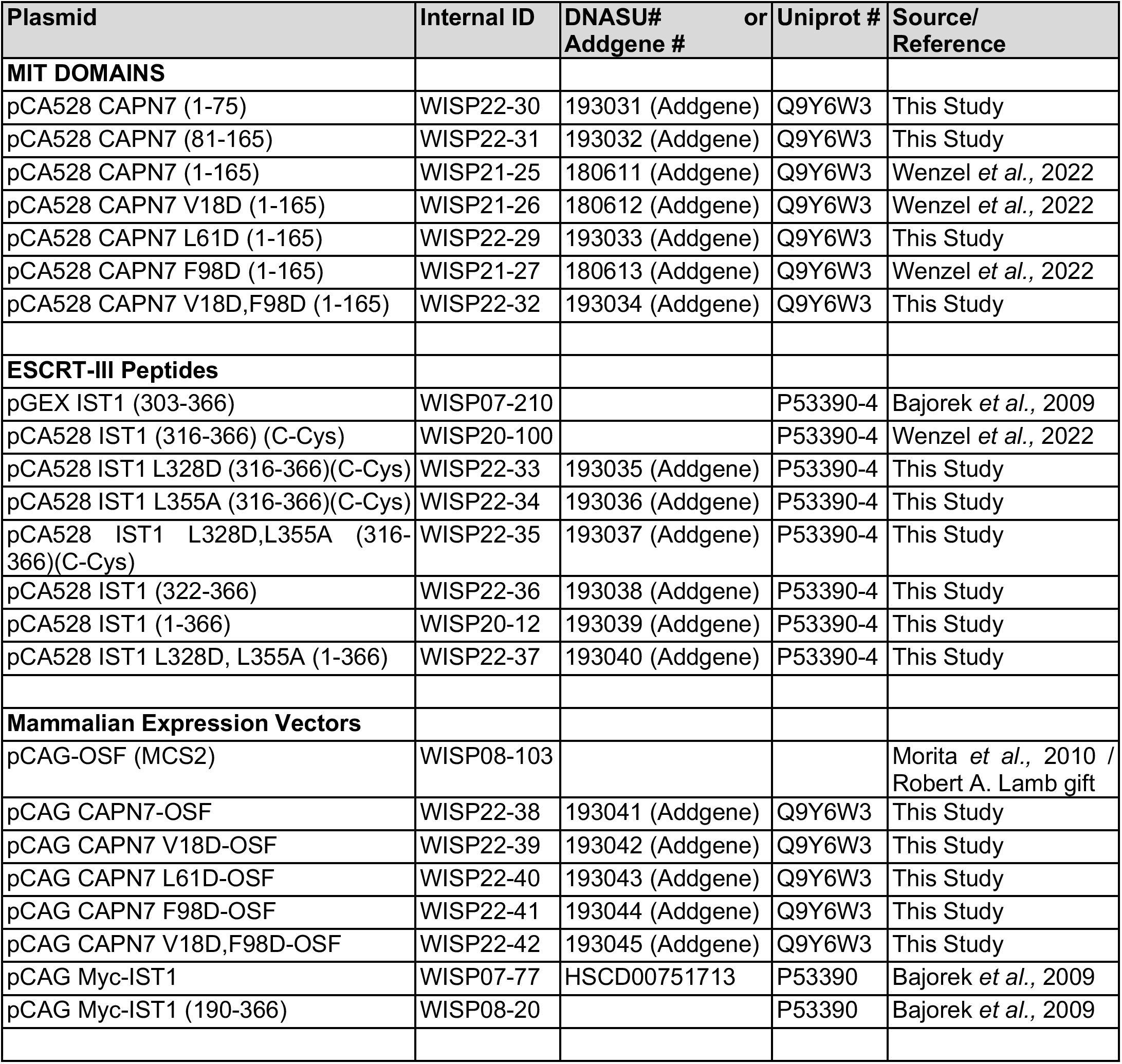
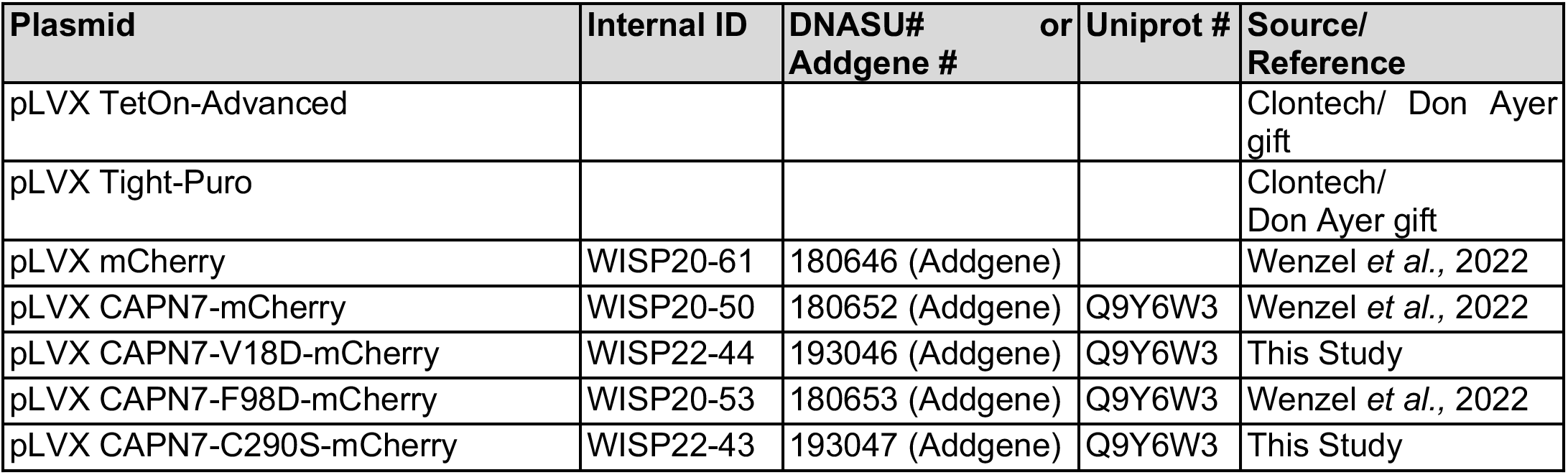
Plasmids.

**Supplemental Table 3.**
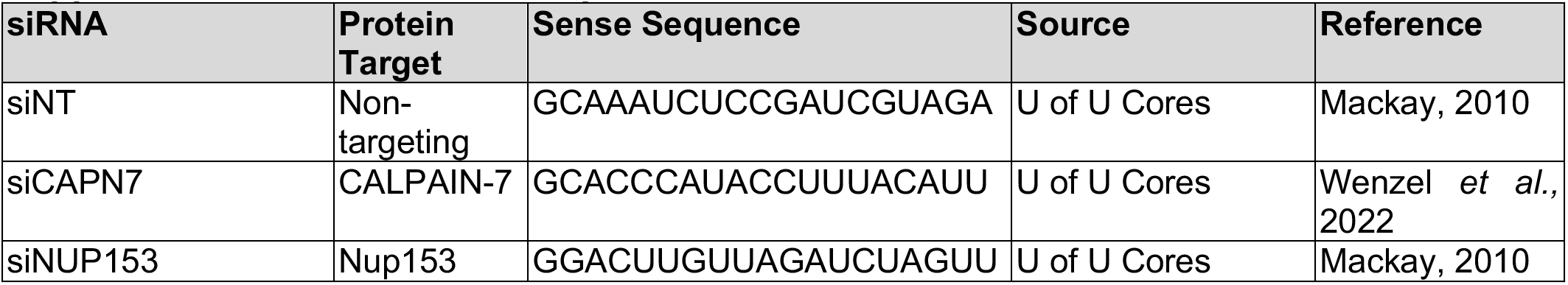
siRNA Sequences.

**Supplemental Table 4.** Antibodies.

| Target | Host Organism | Source | Product Number | Application<sup>1</sup> | Dilution Factor |
| --- | --- | --- | --- | --- | --- |
| CAPN7 | Rabbit | Proteintech | 26985-1-AP | IF<br>WB | 1:500<br>1:4,000 |
| GAPDH | Mouse | Millipore | MAB374 | WB | 1:20,000 |
| IST1 | Rabbit | Sundquist Lab/<br>Covance | UT560 | IF | 1:1,000 |
| mCherry | Rabbit | Abcam | ab167453 | IF | 1:1,000 |
| RFP (mCherry) | Rat | ChromoTek | 5F8 | IF | 1:500 |
| RFP (mCherry) | Mouse | ChromoTek | 6G6 | WB | 1:1,000 |
| NUP153 (SA1) | Mouse | Brian Burke,<br>Singapore | N/A | WB | 1:50 |
| NUP50 | Rabbit | Mackay, et al., 2010 | N/A | WB | 1:2,500 |
| α-TUBULIN (DM1A) | Mouse | Cell Signaling<br>Technology | 3873S | IF | 1:2,000 |
| α-TUBULIN | Chicken | Synaptic Systems | 302 206 | IF | 1:1,000 |
| Myc | Mouse | EMD Millipore | Clone 4A6 | WB | 1:4,000 |
| Flag | Mouse | Sigma | M2 | WB | 1:5,000 |
| IRDye 800 CW,<br>Mouse | Donkey | Licor | 926-32212 | WB | 1:10,000 |
| IRDye 680,<br>Mouse | Donkey | Licor | 926-68072 | WB | 1:10,000 |
| IRDye 800 CW,<br>Rabbit | Donkey | LiCor | 926-32213 | WB | 1:10,000 |
| IRDye 680, Rabbit | Donkey | LiCor | 926-68073 | WB | 1:10,000 |
| DyLight 405<br>AffiniPure, Chicken<br>IgG | Donkey | Jackson<br>Immunoresearch | 703-475-155 | IF | 1:500 |
| Alexa Fluor 594<br>AffiniPure, Rat IgG | Donkey | Jackson<br>Immunoresearch | 712-585-150 | IF | 1:500 |
| Alexa Fluor Plus 488,<br>Rabbit IgG | Donkey | Thermo Fisher | A-32790 | IF | 1:1,000 |
| Alexa Fluor Plus 488,<br>Mouse IgG | Donkey | Thermo Fisher | A-32766 | IF | 1:1,000 |
| Alexa Fluor Plus 594,<br>Rabbit IgG | Donkey | Thermo Fisher | A-32754 | IF | 1:1,000 |

<sup>1</sup>IF = Immunofluorescence; WB = Western Blot

**Figure 1 - figure supplement 1.**
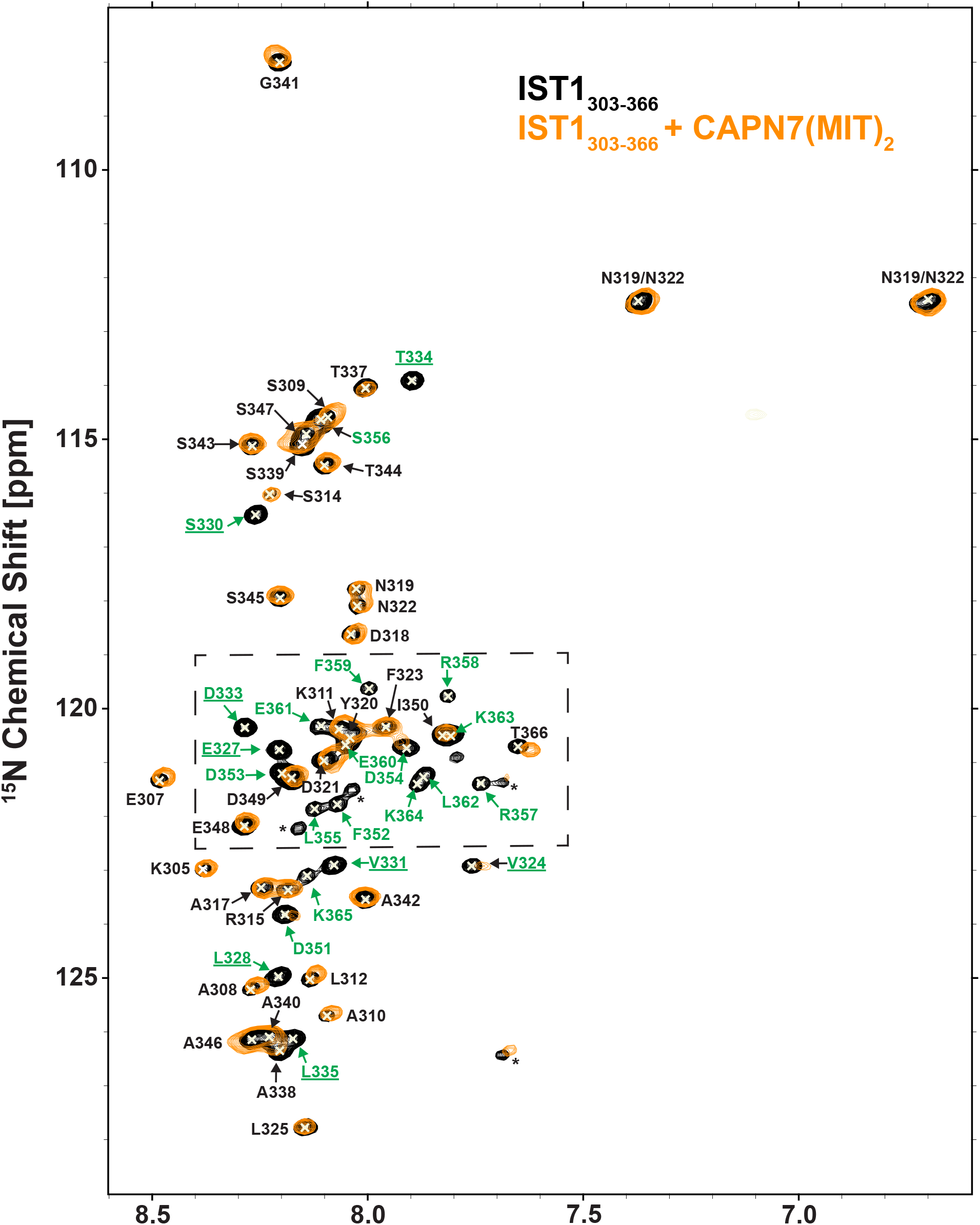
NMR spectra of free and CAPN7(MIT)_2_ bound ^15^N-labeled IST1_303-366_. Overlaid 2D [^15^N,^1^H] HSQC spectra from uniformly ^15^N-labeled IST1_303-366_ in the absence (black contours) and presence (orange contours) of saturating levels of CAPN7(MIT)_2_. Boxed region shows the subset of the spectra shown in Fig. 1B. Asterisks denote unassigned NH signals arising from proline cis/trans isomerization or degradation products.

**Figure 1 – figure supplement 2.**
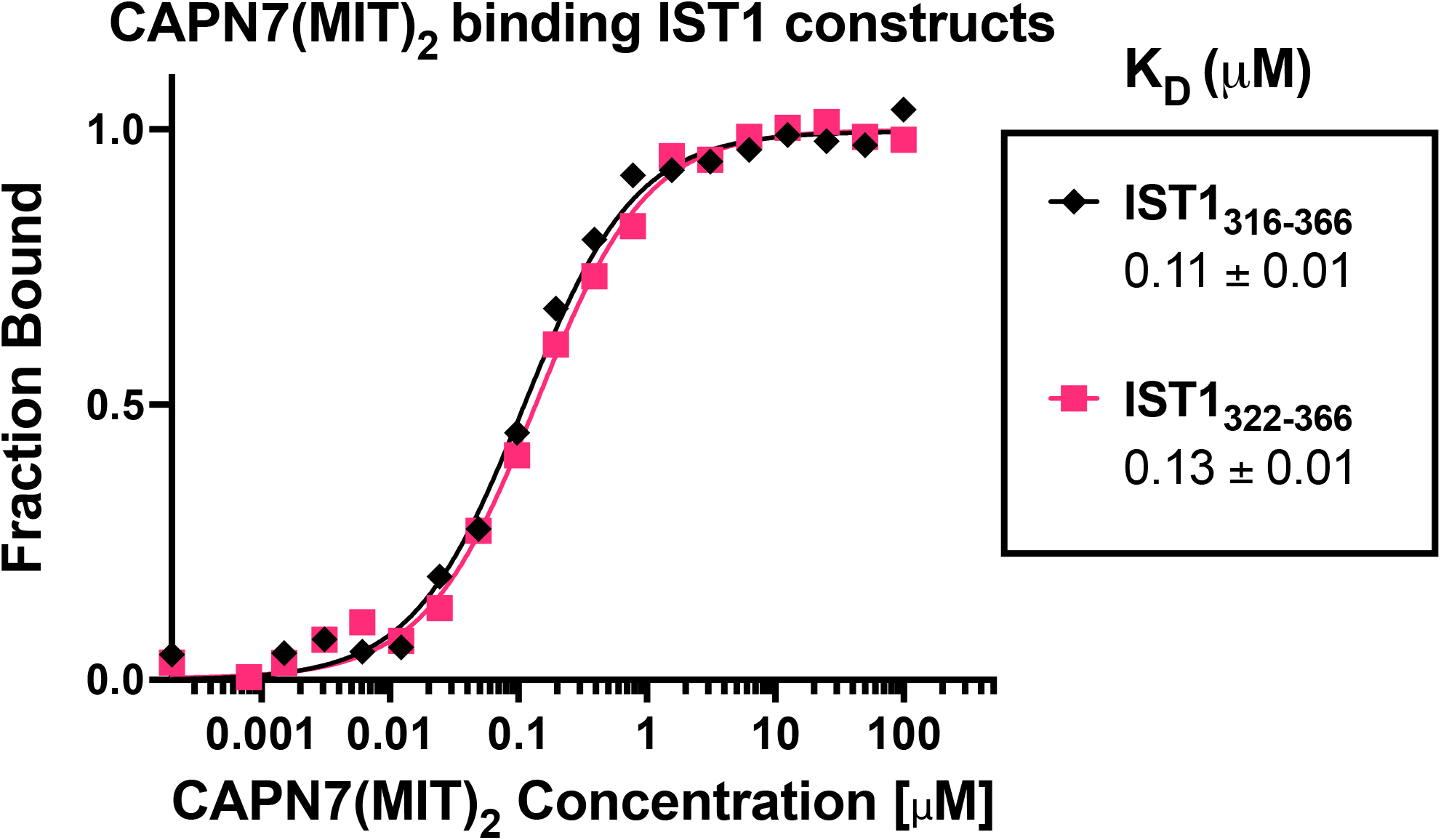
CAPN7(MIT)_2_ binds equally well to IST1_316-366_ and the minimal binding construct IST1_322-366_. Fluorescence polarization anisotropy binding isotherms showing CAPN7(MIT)_2_ binding to fluorescently-labelled peptides spanning IST1_316-366_ or the minimal construct defined by NMR chemical shift mapping (IST1_322-366_). Isotherm data points and dissociation constants are means from three independent experiments ± standard error.

**Figure 2 – figure supplement 1.**
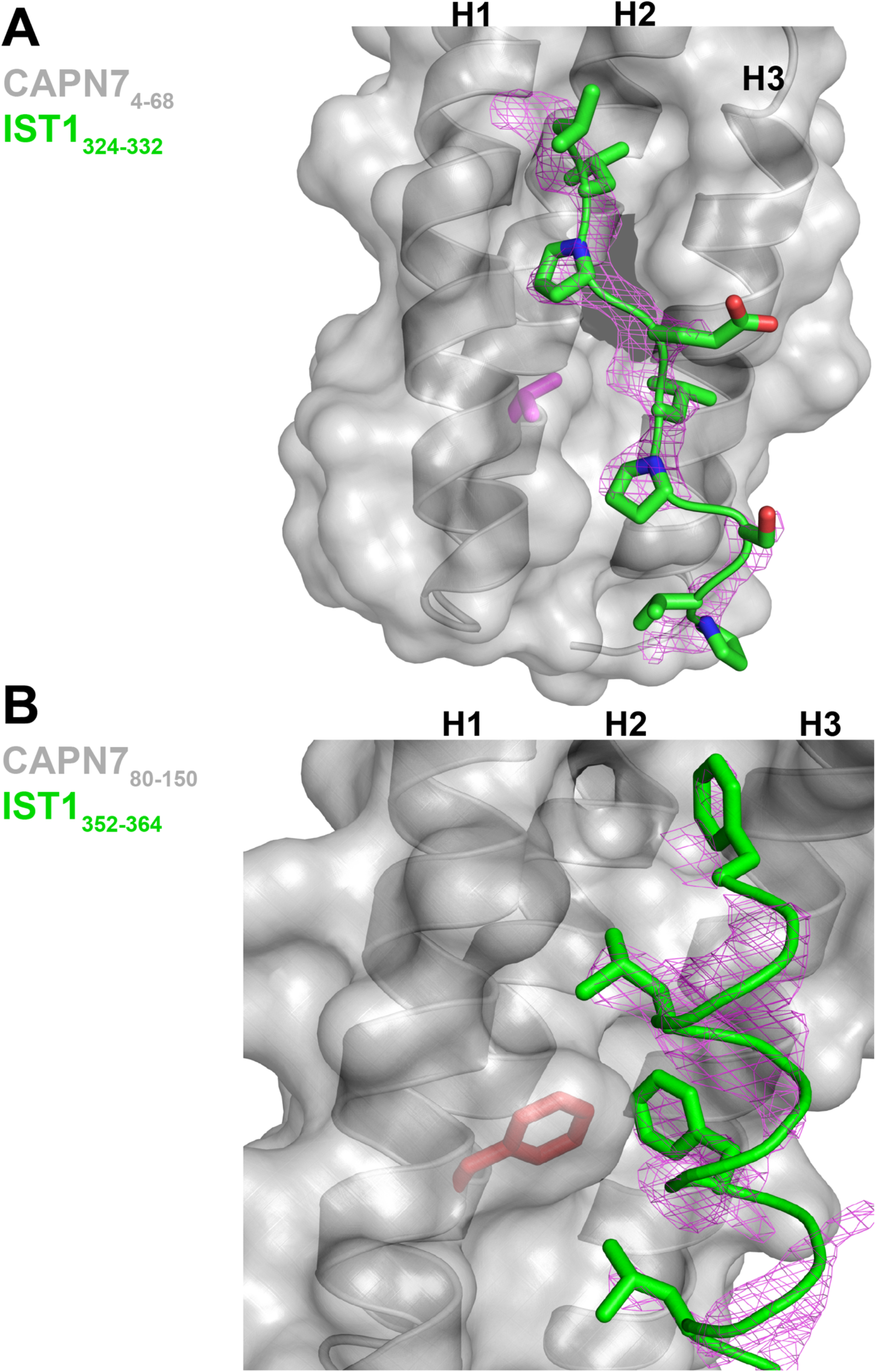
Unbiased electron density omit map for IST1_322-366_. A) IST1_324-332_ Fo-Fc omit map (magenta, contoured at 2.0 σ) overlaid with the CAPN7_4-68_-IST1_324-332_ complex. The CAPN7 Val18 residue is highlighted in magenta. B) IST1_349-364_ Fo-Fc omit map (magenta, contoured at 2.0 σ) overlaid with the CAPN7_80-150_–IST1_349-364_ complex. The CAPN7 Phe98 residue is highlighted in red.

**Figure 2 – figure supplement 2.**
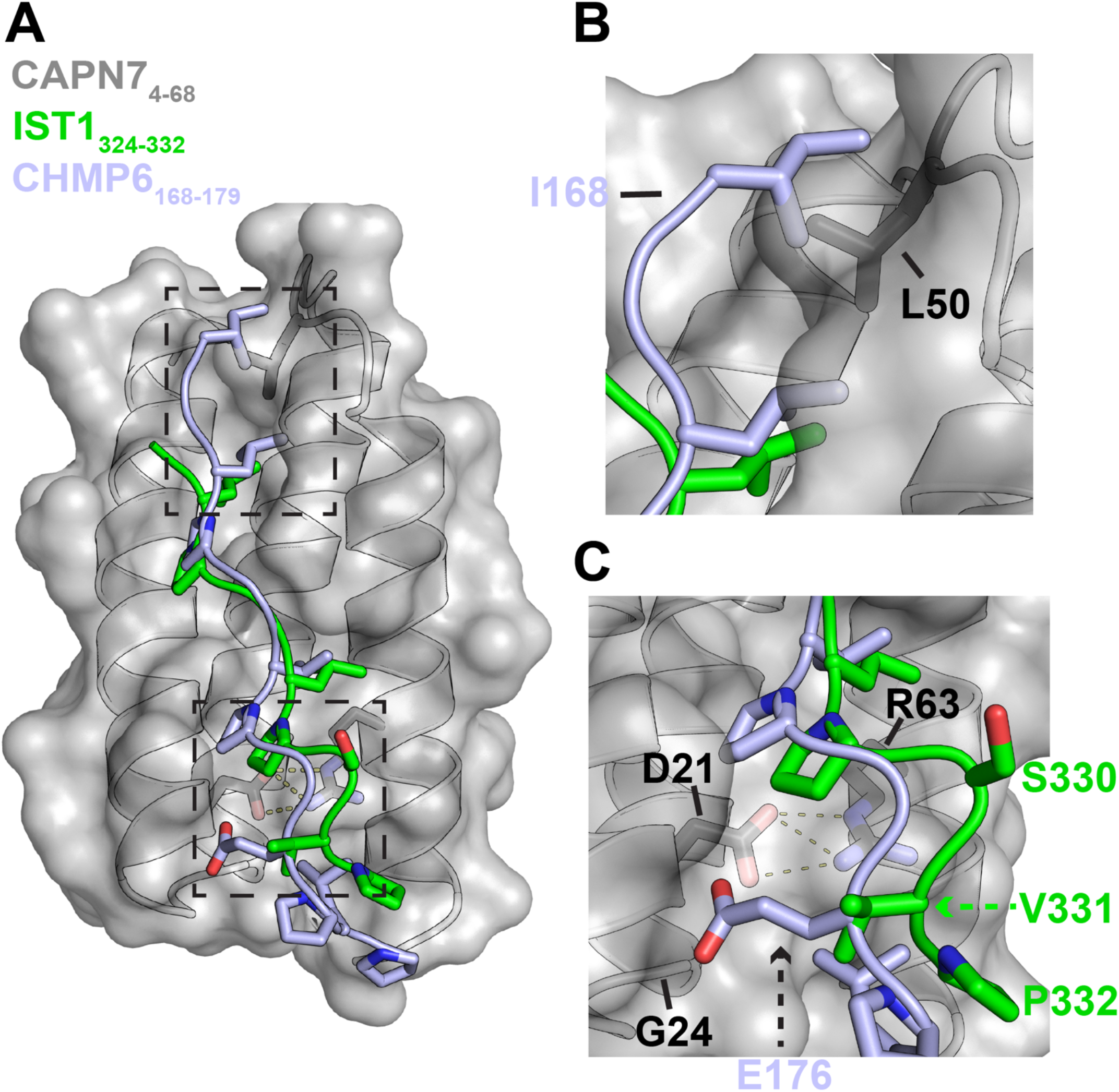
Alignment of IST1_325-336_ and CHMP6_168-179_ in the CAPN7_4-68_ binding groove. A) Structural alignment of the CAPN7_4-68_–IST1_324-332_ complex (gray and green) with CHMP6_168-179_ (light blue) from the VPS4A_3-75_-CHMP6_168-179_ complex (PDB 2K3W). CAPN7_4-68_ is shown as a surface representation in grey with relevant sidechains shown in black. Note the similarities in the core binding region of IST1_324-332_ and CHMP6_168-179_. Dashed boxes show regions magnified to highlight the sequence divergence of IST1_324-332_ and CHMP6_168-179_ in panels B) and C). B) Zoomed view of the N-terminal region of CHMP6_168-179_ to highlight the clash between Ile168 and Leu50 of CAPN7. IST1_324-332_ has no modeled residues in this region due to a lack of interpretable electron density. C) Zoomed view of the C-terminal region CHMP6_168-179_ to highlight Glu176 in the hydrophobic groove of CAPN7. CAPN7 has Gly24 instead of Lys23 of VPS4A, which normally stabilizes CHMP6 Glu176 via hydrogen bonding.

**Figure 2 – figure supplement 3.**
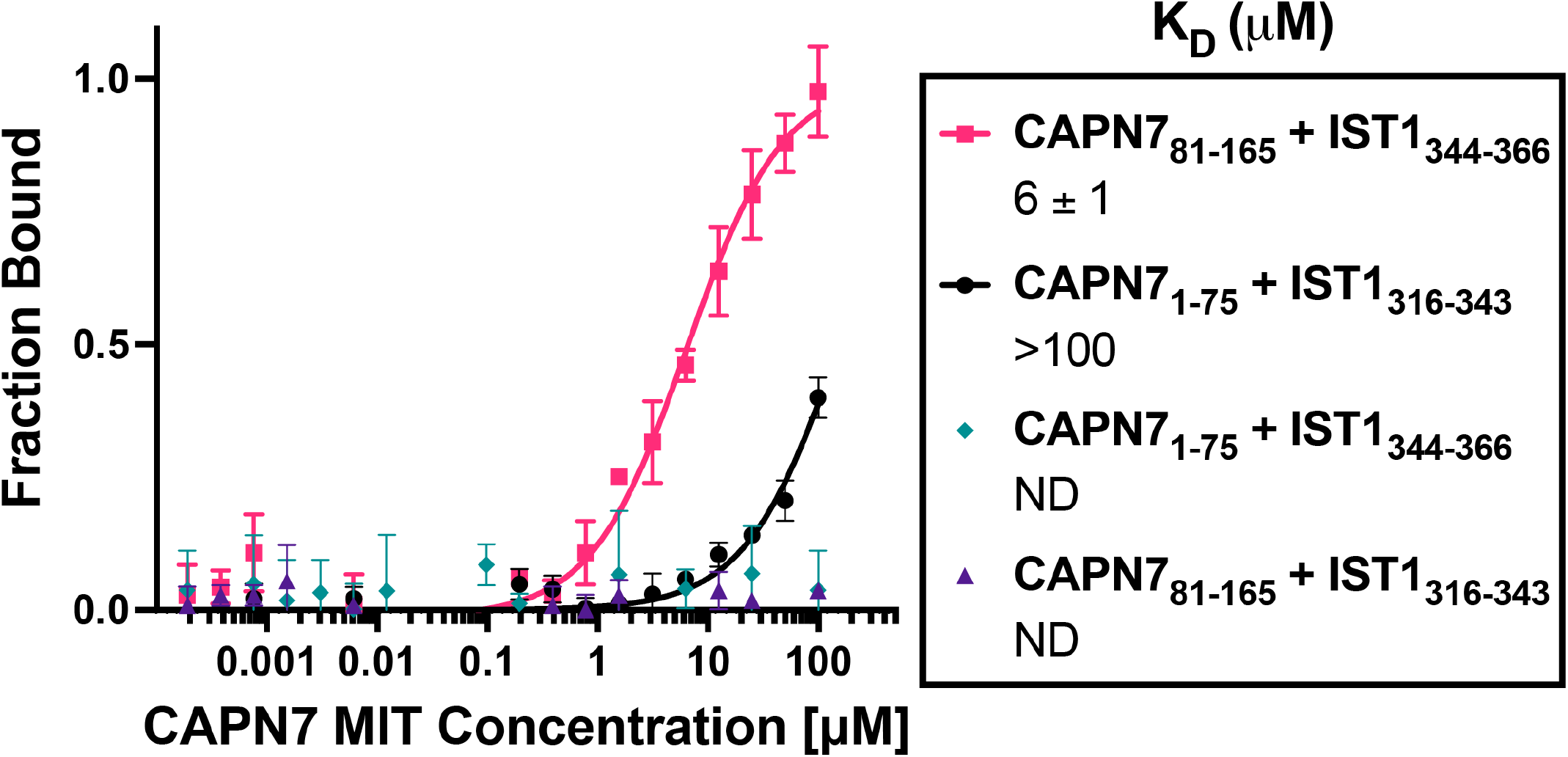
Binding isotherms for individual CAPN7 MIT domains binding to individual IST1 MIM elements. Fluorescence polarization anisotropy binding isotherms show individual CAPN7 MIT domains binding to individual fluorescently-labelled IST1 peptides spanning the N-terminal (IST1_316-343_) or C-terminal (IST1_344-366_) MIM elements. Isotherm data points and dissociation constants are means from three independent experiments ± standard error. ND: Not Determined (see Materials and Methods).

**Figure 3 – figure supplement 1.**
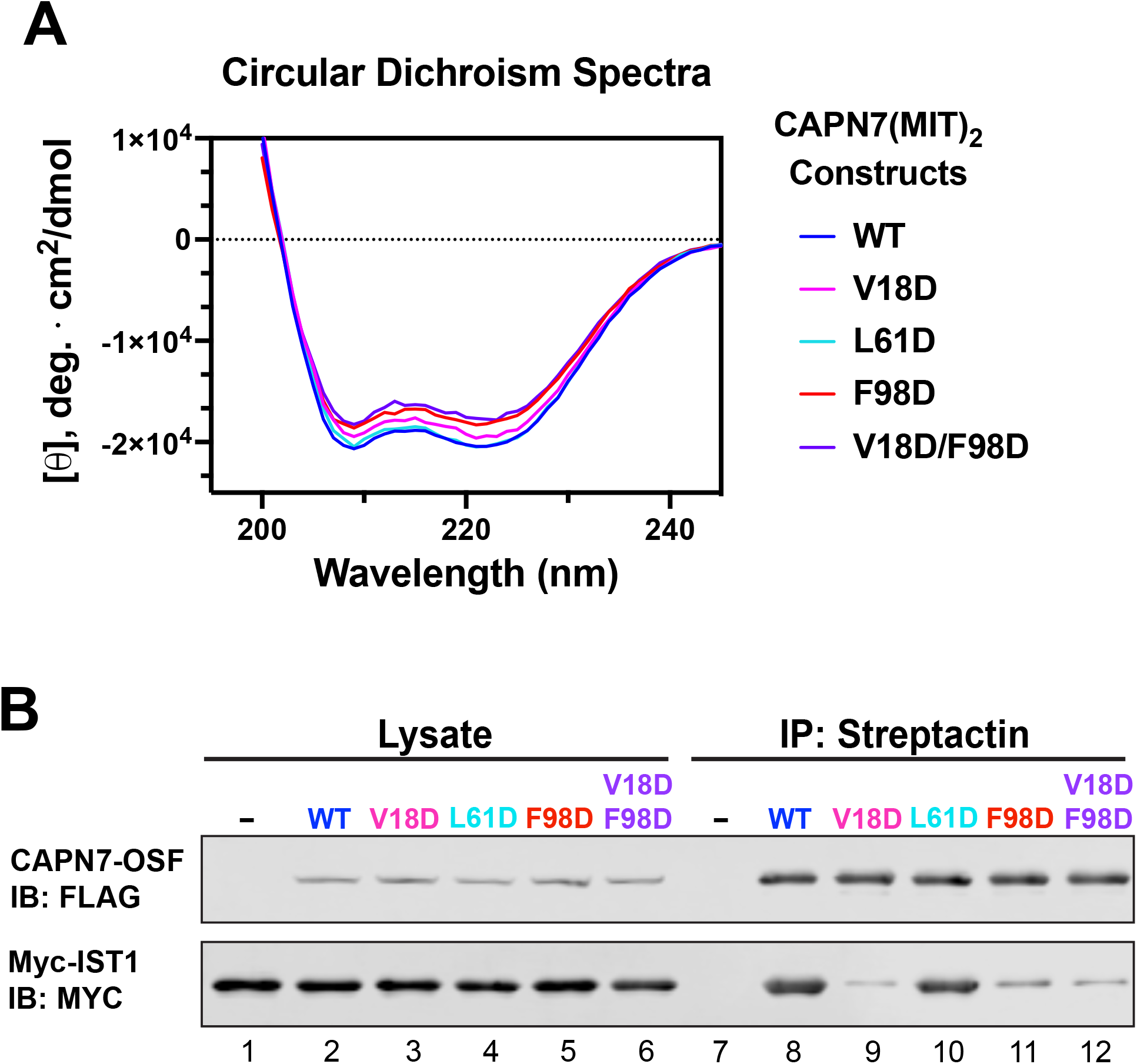
Circular dichroism of CAPN7(MIT)_2_ recombinant proteins and Co-IP of full-length CAPN7 and full-length IST1 from cells. A) Circular dichroism spectra of purified, recombinant CAPN7(MIT)_2_ constructs with indicated point mutations. Spectra are displayed as the mean of triplicate measurements. B) Co-immunoprecipitation of full-length Myc-IST1 and the indicated full-length CAPN7-OSF constructs from extracts of transfected HEK293T cells. Note that the full-length Myc-IST1 showed modest levels of background binding.

**Figure 3 – figure supplement 3.**
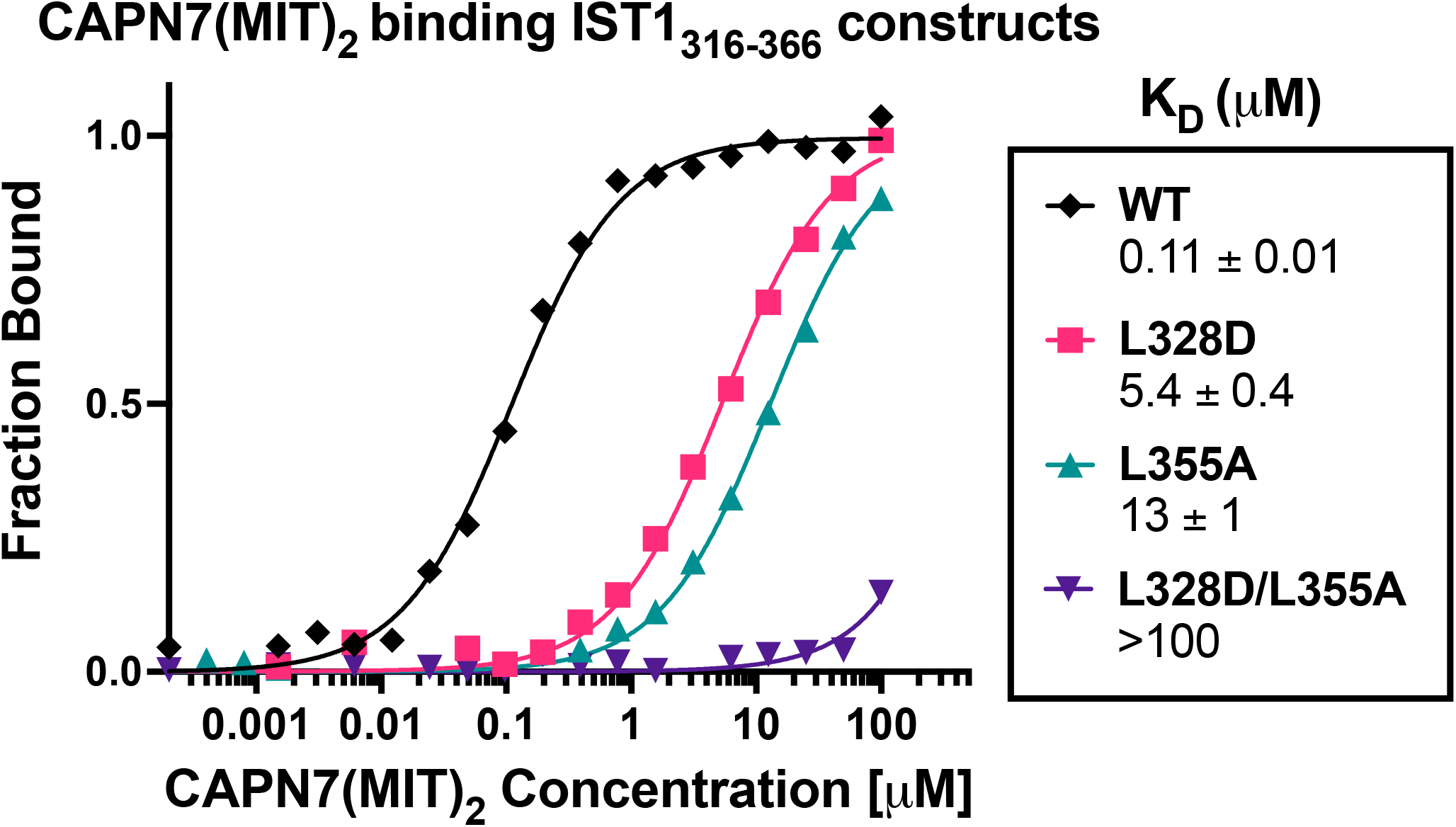
CAPN7(MIT)_2_ binding to IST1_316-366_ is diminished by mutations in either IST1 MIM element. Fluorescence polarization anisotropy binding isotherms showing CAPN7(MIT)_2_ binding to a fluorescently-labelled wt IST1 peptide spanning both MIM elements (IST1_316-366_), or to IST_316-366_ peptides with mutations in either the N-terminal (L328D), C-terminal (L355A) or both (L328D/L355A) MIM elements. Isotherm data points and dissociation constants are means from three independent experiments ± standard error.

**Figure 4 - figure supplement 1.**
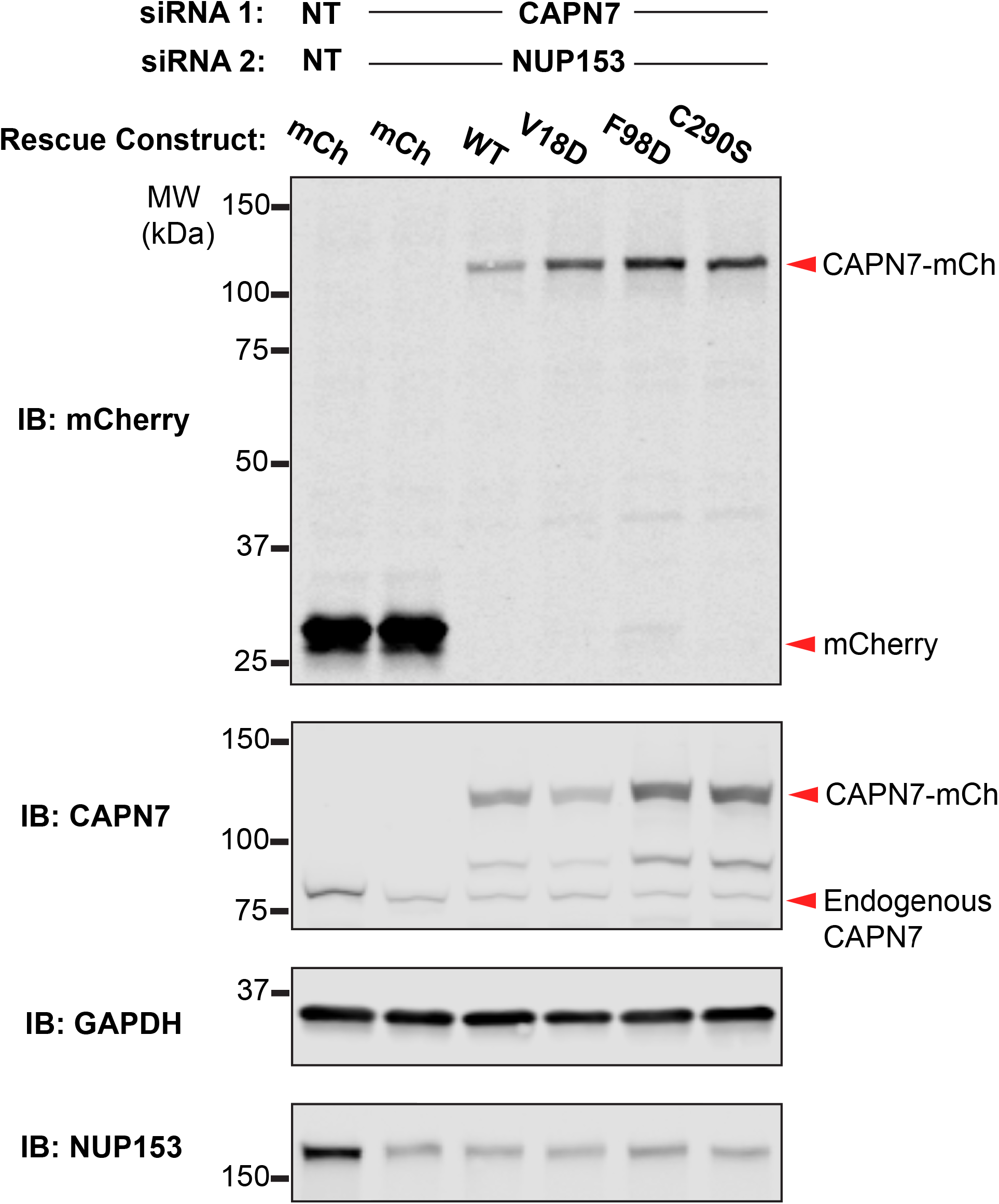
Confirmation of rescue construct expression and siRNA knockdown efficiency. A) Western blot analyses showing CAPN7-mCherry protein expression levels and knockdown efficiencies of endogenous CAPN7 and Nup153 in the experiments shown in Figure 4A and B. siNT is a non-targeting sequence (see materials and methods).

**Figure 5 – figure supplement 1.**
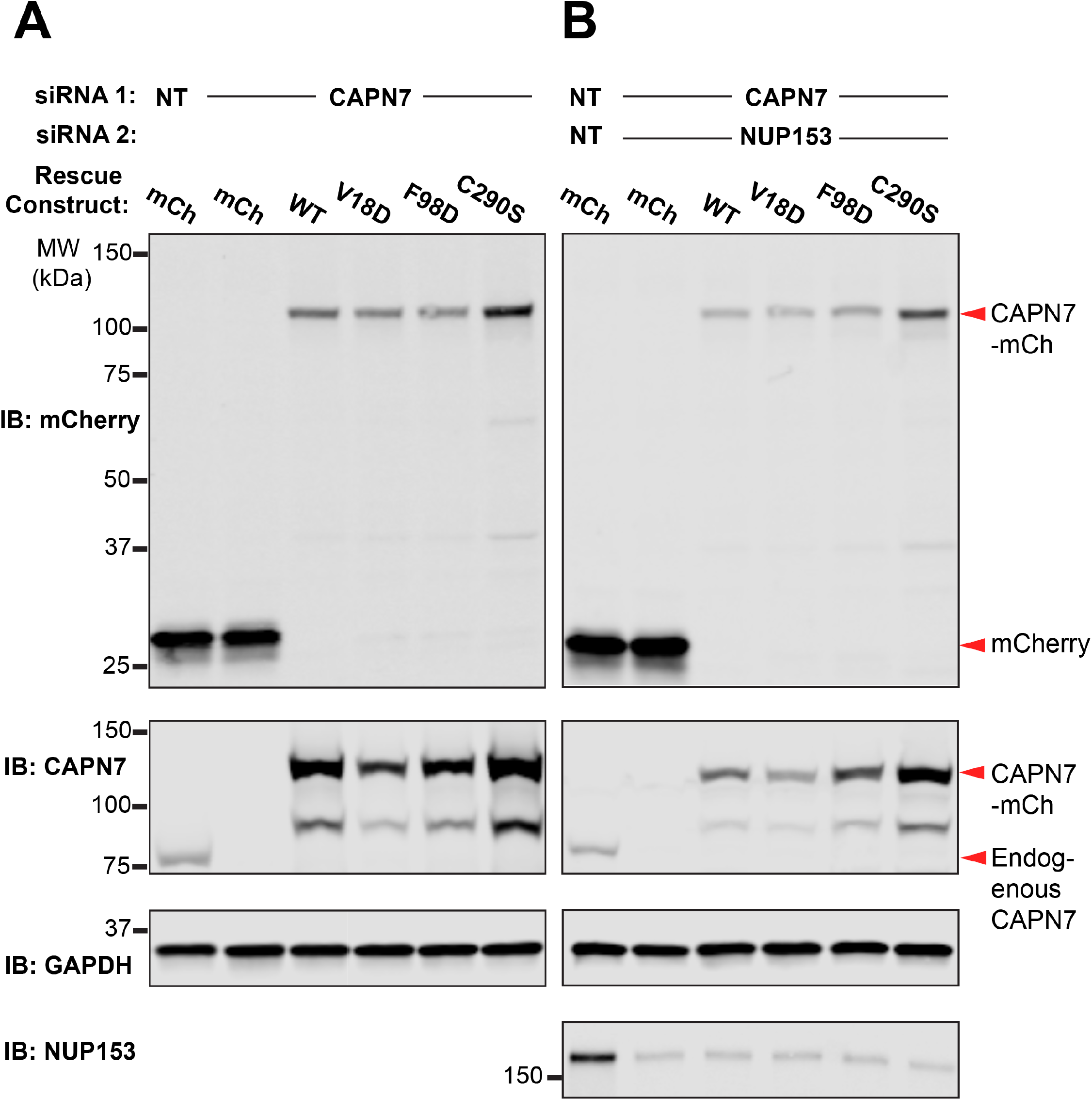
Confirmation of rescue construct expression and siRNA knockdown efficiency. (A, B) Western blot analyses showing mCherry-CAPN7 protein expression levels and knockdown efficiencies of endogenous CAPN7 and Nup153 in the experiments shown in Figure 5A and 5B, respectively.

## Notes

### Competing Interest Statement

The authors have declared no competing interest.

### Summary of Updates

Added Figure 1 - figure supplement 2 and Figure 3 - figure supplement 2.

## References

Agromayor, M., Carlton, J. G., Phelan, J. P., Matthews, D. R., Carlin, L. M., Ameer-Beg, S., Bowers, K., & Martin-Serrano, J. (2009). Essential role of hIST1 in cytokinesis. Mol Biol Cell, 20(5), 1374–1387. doi:10.1091/mbc.E08-05-0474

Azad, K., Guilligay, D., Boscheron, C., Maity, S., De Franceschi, N., Sulbaran, G., Effantin, G., Wang, H., Kleman, J.-P., Bassereau, P., Schoehn, G., Roos, W. H., Desfosses, A., & Weissenhorn, W. (2022). Structural basis of CHMP2A-CHMP3 ESCRT-III polymer assembly and membrane cleavage. bioRxiv, 2022.2004.2012.487901. doi:10.1101/2022.04.12.487901

Bajorek, M., Schubert, H. L., McCullough, J., Langelier, C., Eckert, D. M., Stubblefield, W. M., Uter, N. T., Myszka, D. G., Hill, C. P., & Sundquist, W. I. (2009). Structural basis for ESCRT-III protein autoinhibition. Nat Struct Mol Biol, 16(7), 754–762. doi:10.1038/nsmb.1621

Bunkoczi, G., Echols, N., McCoy, A. J., Oeffner, R. D., Adams, P. D., & Read, R. J. (2013). Phaser.MRage: automated molecular replacement. Acta Crystallogr D Biol Crystallogr, 69(Pt 11), 2276–2286. doi:10.1107/S0907444913022750

Caballe, A., Wenzel, D. M., Agromayor, M., Alam, S. L., Skalicky, J. J., Kloc, M., Carlton, J. G., Labrador, L., Sundquist, W. I., & Martin-Serrano, J. (2015). ULK3 regulates cytokinetic abscission by phosphorylating ESCRT-III proteins. Elife, 4, e06547. doi:10.7554/eLife.06547

Capalbo, L., Mela, I., Abad, M. A., Jeyaprakash, A. A., Edwardson, J. M., & D’Avino, P. P. (2016). Coordinated regulation of the ESCRT-III component CHMP4C by the chromosomal passenger complex and centralspindlin during cytokinesis. Open Biol, 6(10). doi:10.1098/rsob.160248

Capalbo, L., Montembault, E., Takeda, T., Bassi, Z. I., Glover, D. M., & D’Avino, P. P. (2012). The chromosomal passenger complex controls the function of endosomal sorting complex required for transport-III Snf7 proteins during cytokinesis. Open Biol, 2(5), 120070. doi:10.1098/rsob.120070

Carlton, J. G., Caballe, A., Agromayor, M., Kloc, M., & Martin-Serrano, J. (2012). ESCRT-III governs the Aurora B-mediated abscission checkpoint through CHMP4C. Science, 336(6078), 220–225. doi:10.1126/science.1217180

Carlton, J. G., & Martin-Serrano, J. (2007). Parallels between cytokinesis and retroviral budding: a role for the ESCRT machinery. Science, 316(5833), 1908–1912. doi:10.1126/science.1143422

Denison, S. H., Orejas, M., & Arst, H. N., Jr. (1995). Signaling of ambient pH in Aspergillus involves a cysteine protease. J Biol Chem, 270(48), 28519–28522. doi:10.1074/jbc.270.48.28519

Elia, N., Fabrikant, G., Kozlov, M. M., & Lippincott-Schwartz, J. (2012). Computational model of cytokinetic abscission driven by ESCRT-III polymerization and remodeling. Biophys J, 102(10), 2309–2320. doi:10.1016/j.bpj.2012.04.007

Elia, N., Sougrat, R., Spurlin, T. A., Hurley, J. H., & Lippincott-Schwartz, J. (2011). Dynamics of endosomal sorting complex required for transport (ESCRT) machinery during cytokinesis and its role in abscission. Proc Natl Acad Sci USA, 108(12), 4846–4851. doi:10.1073/pnas.1102714108

Emsley, P., & Cowtan, K. (2004). Coot: model-building tools for molecular graphics. Acta Crystallogr D Biol Crystallogr, 60(Pt 12 Pt 1), 2126–2132. doi:10.1107/S0907444904019158

Emsley, P., Lohkamp, B., Scott, W. G., & Cowtan, K. (2010). Features and development of Coot. Acta Crystallogr D Biol Crystallogr, 66(Pt 4), 486–501. doi:10.1107/S0907444910007493

Evans, P. R. (2011). An introduction to data reduction: space-group determination, scaling and intensity statistics. Acta Crystallogr D Biol Crystallogr, 67(Pt 4), 282–292. doi:10.1107/S090744491003982X

Evans, P. R., & Murshudov, G. N. (2013). How good are my data and what is the resolution? Acta Crystallogr D Biol Crystallogr, 69(Pt 7), 1204–1214. doi:10.1107/S0907444913000061

Futai, E., Maeda, T., Sorimachi, H., Kitamoto, K., Ishiura, S., & Suzuki, K. (1999). The protease activity of a calpain-like cysteine protease in Saccharomyces cerevisiae is required for alkaline adaptation and sporulation. Mol Gen Genet, 260(6), 559–568. doi:10.1007/s004380050929

Gibson, D. G., Young, L., Chuang, R. Y., Venter, J. C., Hutchison, C. A., 3rd, & Smith, H. O. (2009). Enzymatic assembly of DNA molecules up to several hundred kilobases. Nat Methods, 6(5), 343–345. doi:10.1038/nmeth.1318

Guizetti, J., Schermelleh, L., Mantler, J., Maar, S., Poser, I., Leonhardt, H., Muller-Reichert, T., & Gerlich, D. W. (2011). Cortical Constriction During Abscission Involves Helices of ESCRT-III-Dependent Filaments. Science, 331(6024), 1616–1620. doi:10.1126/science.1201847

Guo, E. Z., & Xu, Z. (2015). Distinct mechanisms of recognizing endosomal sorting complex required for transport III (ESCRT-III) protein IST1 by different microtubule interacting and trafficking (MIT) domains. J Biol Chem, 290(13), 8396–8408. doi:10.1074/jbc.M114.607903

Hurley, J. H., & Yang, D. (2008). MIT domainia. Dev Cell, 14(1), 6–8. doi:10.1016/j.devcel.2007.12.013

Kabsch, W. (1976). Solution for Best Rotation to Relate 2 Sets of Vectors. Acta Crystallographica Section A, 32(Sep1), 922–923. doi:Doi 10.1107/S0567739476001873

Kabsch, W. (2010). Integration, scaling, space-group assignment and post-refinement. Acta Crystallogr D Biol Crystallogr, 66(Pt 2), 133–144. doi:10.1107/S0907444909047374

Kabsch, W. (2010). Xds. Acta Crystallogr D Biol Crystallogr, 66(Pt 2), 125–132. doi:10.1107/S0907444909047337

Kang, N., Shan, H., Wang, J., Mei, J., Jiang, Y., Zhou, J., Huang, C., Zhang, H., Zhang, M., Zhen, X., Yan, G., & Sun, H. (2022). Calpain7 negatively regulates human endometrial stromal cell decidualization in EMs by promoting FoxO1 nuclear exclusion via hydrolyzing AKT1. Biol Reprod, 106(6), 1112–1125. doi:10.1093/biolre/ioac041

Kieffer, C., Skalicky, J. J., Morita, E., De Domenico, I., Ward, D. M., Kaplan, J., & Sundquist, W. I. (2008). Two distinct modes of ESCRT-III recognition are required for VPS4 functions in lysosomal protein targeting and HIV-1 budding. Dev Cell, 15(1), 62–73. doi:10.1016/j.devcel.2008.05.014

Kojima, R., Obita, T., Onoue, K., & Mizuguchi, M. (2016). Structural Fine-Tuning of MIT-Interacting Motif 2 (MIM2) and Allosteric Regulation of ESCRT-III by Vps4 in Yeast. J Mol Biol, 428(11), 2392–2404. doi:10.1016/j.jmb.2016.04.007

Krissinel, E., & Henrick, K. (2007). Inference of macromolecular assemblies from crystalline state. J Mol Biol, 372(3), 774–797. doi:10.1016/j.jmb.2007.05.022

Laskowski, R. A., & Swindells, M. B. (2011). LigPlot+: multiple ligand-protein interaction diagrams for drug discovery. J Chem Inf Model, 51(10), 2778–2786. doi:10.1021/ci200227u

Liebschner, D., Afonine, P. V., Baker, M. L., Bunkoczi, G., Chen, V. B., Croll, T. I., Hintze, B., Hung, L. W., Jain, S., McCoy, A. J., Moriarty, N. W., Oeffner, R. D., Poon, B. K., Prisant, M. G., Read, R. J., Richardson, J. S., Richardson, D. C., Sammito, M. D., Sobolev, O. V., Stockwell, D. H., Terwilliger, T. C., Urzhumtsev, A. G., Videau, L. L., Williams, C. J., & Adams, P. D. (2019). Macromolecular structure determination using X-rays, neutrons and electrons: recent developments in Phenix. Acta Crystallogr D Struct Biol, 75(Pt 10), 861–877. doi:10.1107/S2059798319011471

Liu, H., Jiang, Y., Jin, X., Zhu, L., Shen, X., Zhang, Q., Wang, B., Wang, J., Hu, Y., Yan, G., & Sun, H. (2013). CAPN 7 promotes the migration and invasion of human endometrial stromal cell by regulating matrix metalloproteinase 2 activity. Reprod Biol Endocrinol, 11, 64. doi:10.1186/1477-7827-11-64

Mackay, D. R., Makise, M., & Ullman, K. S. (2010). Defects in nuclear pore assembly lead to activation of an Aurora B-mediated abscission checkpoint. J Cell Biol, 191(5), 923–931. doi:10.1083/jcb.201007124

Maemoto, Y., Kiso, S., Shibata, H., & Maki, M. (2013). Analysis of limited proteolytic activity of calpain-7 using non-physiological substrates in mammalian cells. FEBS J, 280(11), 2594–2607. doi:10.1111/febs.12243

Maemoto, Y., Ono, Y., Kiso, S., Shibata, H., Takahara, T., Sorimachi, H., & Maki, M. (2014). Involvement of calpain-7 in epidermal growth factor receptor degradation via the endosomal sorting pathway. FEBS J, 281(16), 3642–3655. doi:10.1111/febs.12886

Mayya, V., Lundgren, D. H., Hwang, S. I., Rezaul, K., Wu, L., Eng, J. K., Rodionov, V., & Han, D. K. (2009). Quantitative phosphoproteomic analysis of T cell receptor signaling reveals system-wide modulation of protein-protein interactions. Sci Signal, 2(84), ra46. doi:10.1126/scisignal.2000007

Mierzwa, B. E., Chiaruttini, N., Redondo-Morata, L., von Filseck, J. M., Konig, J., Larios, J., Poser, I., Muller-Reichert, T., Scheuring, S., Roux, A., & Gerlich, D. W. (2017). Dynamic subunit turnover in ESCRT-III assemblies is regulated by Vps4 to mediate membrane remodelling during cytokinesis. Nat Cell Biol, 19(7), 787–798. doi:10.1038/ncb3559

Morita, E., Colf, L. A., Karren, M. A., Sandrin, V., Rodesch, C. K., & Sundquist, W. I. (2010). Human ESCRT-III and VPS4 proteins are required for centrosome and spindle maintenance. Proc Natl Acad Sci U S A, 107(29), 12889–12894. doi:10.1073/pnas.1005938107

Morita, E., Sandrin, V., Chung, H. Y., Morham, S. G., Gygi, S. P., Rodesch, C. K., & Sundquist, W. I. (2007). Human ESCRT and ALIX proteins interact with proteins of the midbody and function in cytokinesis. EMBO J, 26(19), 4215–4227. doi:10.1038/sj.emboj.7601850

Nguyen, H. C., Talledge, N., McCullough, J., Sharma, A., Moss, F. R., 3rd, Iwasa, J. H., Vershinin, M. D., Sundquist, W. I., & Frost, A. (2020). Membrane constriction and thinning by sequential ESCRT-III polymerization. Nat Struct Mol Biol, 27(4), 392–399. doi:10.1038/s41594-020-0404-x

Obita, T., Saksena, S., Ghazi-Tabatabai, S., Gill, D. J., Perisic, O., Emr, S. D., & Williams, R. L. (2007). Structural basis for selective recognition of ESCRT-III by the AAA ATPase Vps4. Nature, 449(7163), 735–739. doi:10.1038/nature06171

Orejas, M., Espeso, E. A., Tilburn, J., Sarkar, S., Arst, H. N., & Penalva, M. A. (1995). Activation of the Aspergillus Pacc Transcription Factor in Response to Alkaline Ambient Ph Requires Proteolysis of the Carboxy-Terminal Moiety. Genes & Development, 9(13), 1622–1632. doi:DOI 10.1101/gad.9.13.1622

Osako, Y., Maemoto, Y., Tanaka, R., Suzuki, H., Shibata, H., & Maki, M. (2010). Autolytic activity of human calpain 7 is enhanced by ESCRT-III-related protein IST1 through MIT-MIM interaction. FEBS J, 277(21), 4412–4426. doi:10.1111/j.1742-4658.2010.07822.x

Penalva, M. A., Lucena-Agell, D., & Arst, H. N., Jr. (2014). Liaison alcaline: Pals entice non-endosomal ESCRTs to the plasma membrane for pH signaling. Curr Opin Microbiol, 22, 49–59. doi:10.1016/j.mib.2014.09.005

Pfitzner, A. K., Mercier, V., Jiang, X., Moser von Filseck, J., Baum, B., Saric, A., & Roux, A. (2020). An ESCRT-III Polymerization Sequence Drives Membrane Deformation and Fission. Cell, 182(5), 1140-1155 e1118. doi:10.1016/j.cell.2020.07.021

Pfitzner, A. K., Moser von Filseck, J., & Roux, A. (2021). Principles of membrane remodeling by dynamic ESCRT-III polymers. Trends Cell Biol, 31(10), 856–868. doi:10.1016/j.tcb.2021.04.005

Rodriguez-Galan, O., Galindo, A., Hervas-Aguilar, A., Arst, H. N., Jr., & Penalva, M. A. (2009). Physiological involvement in pH signaling of Vps24-mediated recruitment of Aspergillus PalB cysteine protease to ESCRT-III. J Biol Chem, 284(7), 4404–4412. doi:10.1074/jbc.M808645200

Samson, R. Y., Obita, T., Freund, S. M., Williams, R. L., & Bell, S. D. (2008). A role for the ESCRT system in cell division in archaea. Science, 322(5908), 1710–1713. doi:10.1126/science.1165322

Scott, A., Gaspar, J., Stuchell-Brereton, M. D., Alam, S. L., Skalicky, J. J., & Sundquist, W. I. (2005). Structure and ESCRT-III protein interactions of the MIT domain of human VPS4A. Proc Natl Acad Sci U S A, 102(39), 13813–13818. doi:10.1073/pnas.0502165102

Scourfield, E. J., & Martin-Serrano, J. (2017). Growing functions of the ESCRT machinery in cell biology and viral replication. Biochem Soc Trans, 45(3), 613–634. doi:10.1042/BST20160479

Skalicky, J. J., Arii, J., Wenzel, D. M., Stubblefield, W. M., Katsuyama, A., Uter, N. T., Bajorek, M., Myszka, D. G., & Sundquist, W. I. (2012). Interactions of the human LIP5 regulatory protein with endosomal sorting complexes required for transport. J Biol Chem, 287(52), 43910–43926. doi:10.1074/jbc.M112.417899

Strohacker, L. K., Mackay, D. R., Whitney, M. A., Couldwell, G. C., Sundquist, W. I., & Ullman, K. S. (2021). Identification of abscission checkpoint bodies as structures that regulate ESCRT factors to control abscission timing. Elife, 10. doi:10.7554/eLife.63743

Stuchell-Brereton, M. D., Skalicky, J. J., Kieffer, C., Karren, M. A., Ghaffarian, S., & Sundquist, W. I. (2007). ESCRT-III recognition by VPS4 ATPases. Nature, 449(7163), 740–744. doi:10.1038/nature06172

Studier, F. W. (2005). Protein production by auto-induction in high density shaking cultures. Protein Expr Purif, 41(1), 207–234. doi:10.1016/j.pep.2005.01.016

Talledge, N., McCullough, J., Wenzel, D., Nguyen, H. C., Lalonde, M. S., Bajorek, M., Skalicky, J., Frost, A., & Sundqust, W. I. (2018). The ESCRT-III proteins IST1 and CHMP1B assemble around nucleic acids. bioRxiv, 386532. doi:10.1101/386532

Vild, C. J., & Xu, Z. (2014). Vfa1 binds to the N-terminal microtubule-interacting and trafficking (MIT) domain of Vps4 and stimulates its ATPase activity. J Biol Chem, 289(15), 10378–10386. doi:10.1074/jbc.M113.532960

Vincent, O., Rainbow, L., Tilburn, J., Arst, H. N., Jr., & Penalva, M. A. (2003). YPXL/I is a protein interaction motif recognized by aspergillus PalA and its human homologue, AIP1/Alix. Mol Cell Biol, 23(5), 1647–1655. doi:10.1128/MCB.23.5.1647-1655.2003

Wenzel, D. M., Mackay, D. R., Skalicky, J. J., Paine, E. L., Miller, M. S., Ullman, K. S., & Sundquist, W. I. (2022). Comprehensive analysis of the human ESCRT-III-MIT domain interactome reveals new cofactors for cytokinetic abscission. Elife, 11. doi:10.7554/eLife.77779

Winn, M. D., Ballard, C. C., Cowtan, K. D., Dodson, E. J., Emsley, P., Evans, P. R., Keegan, R. M., Krissinel, E. B., Leslie, A. G., McCoy, A., McNicholas, S. J., Murshudov, G. N., Pannu, N. S., Potterton, E. A., Powell, H. R., Read, R. J., Vagin, A., & Wilson, K. S. (2011). Overview of the CCP4 suite and current developments. Acta Crystallogr D Biol Crystallogr, 67(Pt 4), 235–242. doi:10.1107/S0907444910045749

Xu, W. J., & Mitchell, A. P. (2001). Yeast PalA/AIP1/Alix homolog Rim20p associates with a PEST-like region and is required for its proteolytic cleavage. Journal of Bacteriology, 183(23), 6917–6923. doi:Doi 10.1128/Jb.183.23.6917-6923.2001

Yan, Q., Huang, C., Jiang, Y., Shan, H., Jiang, R., Wang, J., Liu, J., Ding, L., Yan, G., & Sun, H. (2018). Calpain7 impairs embryo implantation by downregulating beta3-integrin expression via degradation of HOXA10. Cell Death Dis, 9(3), 291. doi:10.1038/s41419-018-0317-3

Yang, D., Rismanchi, N., Renvoise, B., Lippincott-Schwartz, J., Blackstone, C., & Hurley, J. H. (2008). Structural basis for midbody targeting of spastin by the ESCRT-III protein CHMP1B. Nat Struct Mol Biol, 15(12), 1278–1286. doi:10.1038/nsmb.1512

Yorikawa, C., Takaya, E., Osako, Y., Tanaka, R., Terasawa, Y., Hamakubo, T., Mochizuki, Y., Iwanari, H., Kodama, T., Maeda, T., Hitomi, K., Shibata, H., & Maki, M. (2008). Human calpain 7/PalBH associates with a subset of ESCRT-III-related proteins in its N-terminal region and partly localizes to endocytic membrane compartments. J Biochem, 143(6), 731–745. doi:10.1093/jb/mvn030

